# Disease-associated variants alter a novel physiological role of hnRNP H2 in neuronal development

**DOI:** 10.64898/2026.10.02.756225

**Authors:** Afonso Lourinho, Marta S. Silva, Emanuel Tahiri, Marco D’Ambrosio, Ana D. Pinto, Saray Calvo Parra, Giovana De Pietro, Guilherme Galvão, Ana B. Caniceiro, Luís Pereira de Almeida, Paulo Pinheiro, Irina S. Moreira, Rui Jorge Nobre, Miranda Mele, Andrzej W. Cwetsch, Carlos B. Duarte, Rui O. Costa

**Affiliations:** Center for Neuroscience and Cell Biology (CNC), University of Coimbra, 3004-504 Coimbra, Portugal; CIBB - Center for Innovative Biomedicine and Biotechnology, University of Coimbra, 3000-548 Coimbra, Portugal; Institute for Interdisciplinary Research, University of Coimbra, 3030-789 Coimbra, Portugal; Doctoral Programme in Experimental Biology and Biomedicine (PDBEB), University of Coimbra, 3004-504 Coimbra, Portugal; Instituto de Biotecnología y Biomedicina (BIOTECMED), Universidad de Valencia, Departamento de Biología Celular y Biología Funcional, 46100 Burjassot, Spain; GeneT - Gene Therapy Center of Excellence Portugal, University of Coimbra, 3004-504 Coimbra, Portugal; Department of Life Sciences, University of Coimbra, 3000-456 Coimbra, Portugal

## Abstract

Heterogeneous nuclear ribonucleoprotein H2 (hnRNP H2) is a member of the hnRNP family of RNA-binding proteins that regulate neuronal function through multiple aspects of RNA metabolism. Mutations in the nuclear localisation signal (NLS) and quasi-RNA recognition motifs (qRRMs) of hnRNP H2 have recently been linked to a severe developmental syndrome, termed *HNRNPH2*-related neurodevelopmental disorder. Despite its emerging importance, the physiological roles of hnRNP H2 in neuronal development and synaptic maturation, and how disease-associated variants alter these processes to drive pathology, remain poorly understood. Here, we show that hnRNP H2 is expressed in hippocampal neurons and localises to the synaptic compartment. Knockdown of hnRNP H2 revealed essential roles of the protein in the regulation of neuronal morphology, dendritic spine maturation, synaptic transmission, and cortical neuronal migration *in vivo*. Disease-associated hnRNP H2 variants altered protein localisation and activity-dependent synaptic recruitment, resulting in defects in neuronal morphology, *in vivo* cortical neuronal migration and laminar orientation, dendritic spine maturation, and synaptic function. Together, these findings establish hnRNP H2 as a critical regulator of neurodevelopment and identify a molecular basis for *HNRNPH2*-related neurodevelopmental disorder.

**Graphical Abstract:** 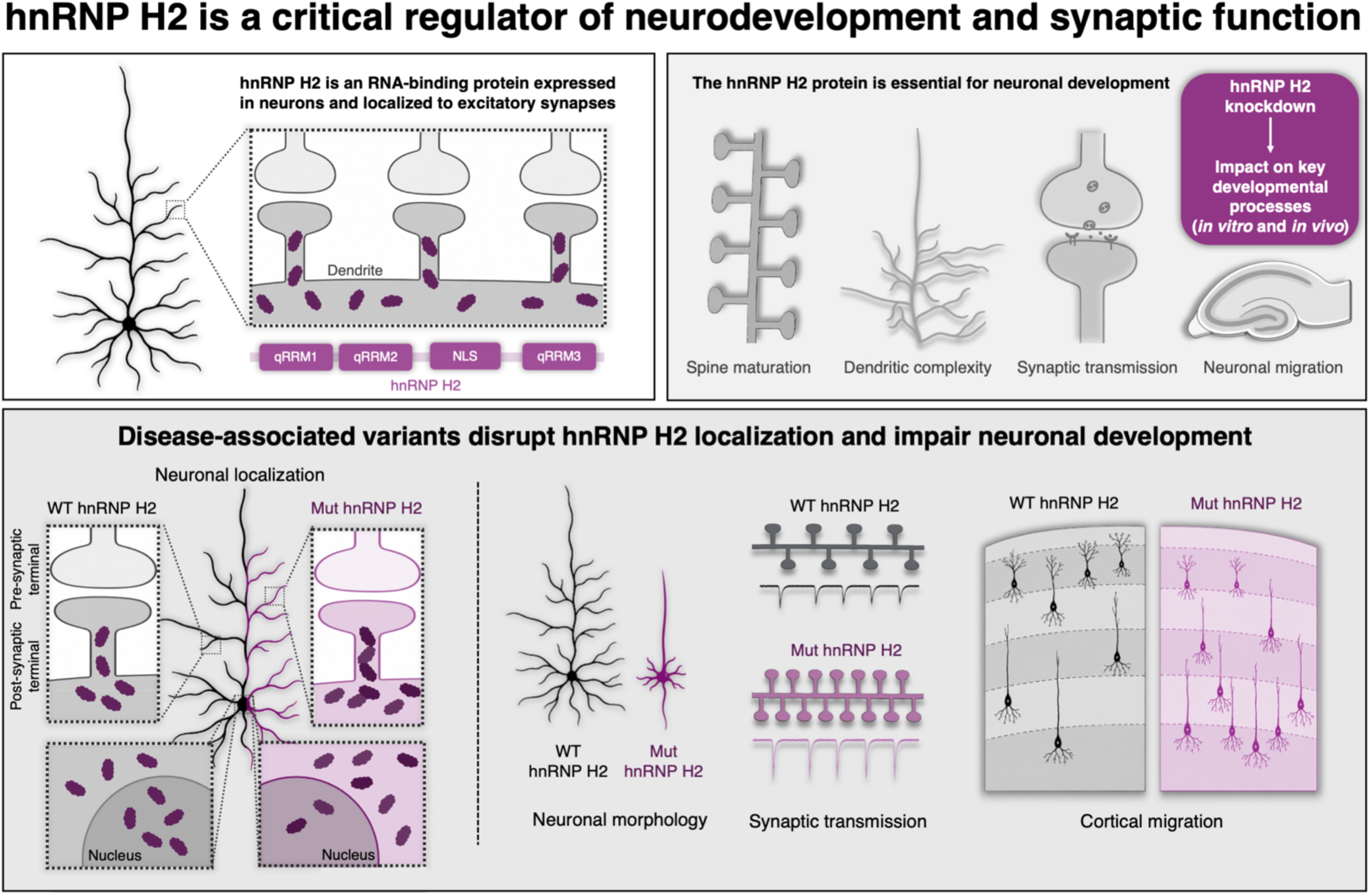

## Introduction

Neurodevelopmental disorders (NDDs) are complex conditions characterized by impairments in cognition, communication, behaviour, and/or psychomotor function. A wide array of genes and genetic mutations have been implicated in NDDs, highlighting the heterogeneous and multifactorial origins of these disorders (Parenti et al., 2020). Intellectual disability (ID), autism spectrum disorder (ASD), attention-deficit/hyperactivity disorder (ADHD) all fall under the NDD umbrella, collectively affecting around 15% of children and adolescents worldwide (Gidziela et al., 2023). Among these, *HNRNPH2*-related neurodevelopmental disorder (*HNRNPH2*-RNDD; OMIM #300986) is a recently described syndrome characterized by developmental delay/intellectual disability, language and motor impairments, growth abnormalities, and musculoskeletal disturbances (Bain et al., 2021). *HNRNPH2*-RNDD is caused by *de novo* mutations in the X-linked *HNRNPH2* gene, which encodes the heterogeneous nuclear ribonucleoprotein H2 (hnRNP H2) (Bain et al., 2016). hnRNP H2 is an RNA-binding protein (RBP) involved in various aspects of RNA metabolism, including alternative splicing, pre-mRNA cleavage, and polyadenylation (Alkan et al., 2006; Bagga et al., 1998; Bain et al., 2016; Stark et al., 2011; Turunen et al., 2013). Despite the well-established roles of hnRNP family proteins in orchestrating synaptic plasticity, axonal integrity and brain development via post-transcriptional RNA regulation (Tilliole et al., 2024; Lourinho et al., 2025), the specific neuronal functions of hnRNP H2 remain largely uncharacterized.

To date, at least 50-70 individuals with *HNRNPH2*-RNDD have been described in scientific literature, although the number of known cases worldwide may be approaching 300 (Garcia et al., 2026; Gillentine et al., 2021). Notably, disease-causing mutations predominantly cluster within the nuclear localisation signal (NLS) and quasi-RNA recognition motifs (qRRMs) of hnRNP H2, domains essential for its nuclear trafficking and RNA-binding properties, respectively (Gillentine et al., 2021; Bain et al., 2016, 2021; Somashekar et al., 2020; White-Brown et al., 2022; Harmsen et al., 2019; Jepsen et al., 2019; Peron et al., 2020; Kreienkamp et al., 2022). Mouse models of *HNRNPH2*-RNDD recapitulate several key clinical features of the disorder, including craniofacial abnormalities, motor deficits, seizures, and cognitive impairments (Korff et al., 2023, 2026). In contrast, *Hnrnph2* knockout mice do not develop these phenotypes, likely due to compensatory upregulation of *Hnrnph1*, a mechanism absent in mutant animals (Korff et al., 2023). Although recent studies have provided important insights into the molecular basis of *HNRNPH2*-RNDD and highlighted potential therapeutic strategies (Korff et al., 2026), the physiological roles of hnRNP H2 during neuronal maturation and the mechanisms by which disease-associated variants perturb neuronal development and function remain unknown. At the molecular level, pathogenic variants appear to exert mutation-specific effects: NLS mutations disrupt nuclear import, resulting in cytoplasmic accumulation of hnRNP H2 and widespread changes in alternative splicing and gene expression, whereas the p.R114W variant within the qRRM domain alters the interaction of hnRNP H2 with nuclear splicing regulators and affects RNA processing (Gonzalez et al., 2023; Korff et al., 2023; Kreienkamp et al., 2022).

A major challenge in understanding the molecular mechanisms underlying neurodevelopmental disorders lies in the limited knowledge of the neuronal functions of many NDD-associated genes. The expression of hnRNP H2 throughout development of the human brain suggests a critical role in neuronal maturation and function, one that may be disrupted by pathogenic mutations in *HNRNPH2* (Kang et al., 2011; Johnson et al., 2009; Pletikos et al., 2014). In this study, we address this major gap by investigating the role of hnRNP H2 in hippocampal neurodevelopment, as well as the morphological, synaptic, and functional consequences of disease-associated hnRNP H2 variants. W demonstrate that hnRNP H2 is expressed in developing hippocampal neurons and is present within the synaptic compartment. We further show that hnRNP H2 regulates dendritic arborization, spine maturation, synaptic transmission, and *in vivo* cortical neuronal migration. Disease-associated hnRNP H2 variants caused protein mislocalisation to dendritic and synaptic compartments and, in a mutation-specific manner, impaired excitatory synaptic transmission while disrupting neuronal morphology, dendritic spine maturation, and cortical neuronal migration and laminar organization. Together, our findings identify hnRNP H2 as a critical mediator of neuronal development and reveal how pathogenic mutations disrupt its function, thereby contributing to the molecular pathology of *HNRNPH2*-RNDD.

## Results

### hnRNP H2 is localised to excitatory synapses in developing hippocampal neurons

The hnRNP H2 protein, encoded by the *HNRNPH2* gene located on chromosome Xq22.1, comprises 449 amino acids and is robustly expressed in the human hippocampus throughout prenatal and postnatal brain development (Kang et al., 2011; Johnson et al., 2009; Pletikos et al., 2014). To investigate the neuronal expression dynamics and subcellular localisation of hnRNP H2, we examined protein levels in primary rat hippocampal neurons across key developmental stages. Immunostaining with an antibody against endogenous hnRNP H2 revealed a pronounced upregulation between days *in vitro* (DIV) 7 and 14, a critical period corresponding to synaptic maturation **(Fig. 1A)**. Prompted by this developmental expression pattern, we next assessed the spatial distribution of hnRNP H2 within hippocampal neurons. While the protein displayed predominant nuclear localisation, we also observed a distinct cytoplasmic pool in the somatodendritic compartment **(Fig. 1B)**. Notably, hnRNP H2 colocalised with Homer1, a marker of the postsynaptic density, suggesting its enrichment at excitatory synapses **(Fig. 1B)**. Confocal imaging revealed hnRNP H2 puncta closely adjacent to Homer1, positioned at the base of dendritic spines, consistent with a postsynaptic localisation **(Fig. 1C, Fig. S1A, Supplementary movie 1)**. To achieve a higher-resolution mapping of synaptic distribution, we employed super-resolution microscopy. hnRNP H2 puncta were consistently found in proximity to both VGluT1 and PSD-95, presynaptic and postsynaptic markers, respectively **(Fig. 1D)**. These findings provide compelling evidence for a previously unrecognized synaptic localisation of the hnRNP H2 protein.

**Figure 1.**
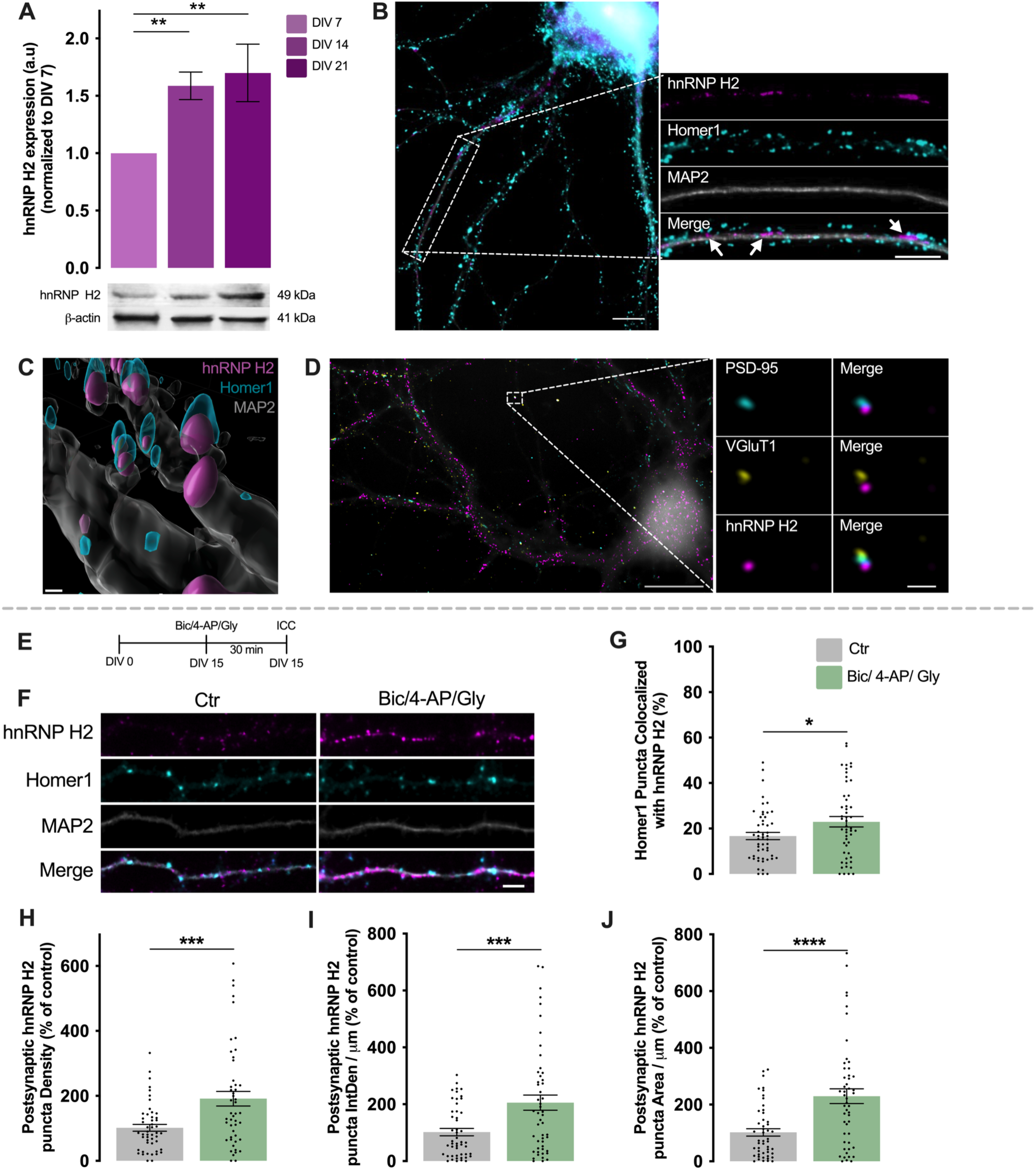
hnRNP H2 localises to excitatory synapses and is recruited in an activity-dependent manner in developing hippocampal neurons. **(A)** Developmental expression of hnRNP H2 in primary cultures of rat hippocampal neurons maintained for 7, 14 and 21 days in vitro (DIV), assessed by immunoblotting. Quantification of hnRNP H2 protein levels normalized to the loading control (β-actin) is shown above. **(B)** Representative confocal images of DIV 21 hippocampal neurons immunostained for hnRNP H2 (magenta), Homer1 (cyan) and MAP2 (grey). Insets show hnRNP H2 puncta localised adjacent to Homer1-positive postsynaptic sites along dendrites (arrows). **(C)** Three-dimensional reconstruction of dendritic segments generated from Airyscan images showing hnRNP H2 localisation at the base of Homer1-positive dendritic spines. The corresponding three-dimensional visualization is provided in Supplementary Movie 1. **(D)** Representative super-resolution image of a hippocampal neuron immunostained for hnRNP H2, VGluT1 and PSD-95. Insets show hnRNP H2 puncta in close apposition to excitatory pre- and postsynaptic markers. Merged images are shown. **(E)** Experimental paradigm for activity-dependent synaptic stimulation. DIV 15 hippocampal neurons were treated with bicuculline (Bic, 50 μM), 4-aminopyridine (4-AP, 2.5 mM) and glycine (Gly, 10 μM) for 30 min before fixation. **(F)** Representative images of dendrites from control and stimulated neurons immunostained for hnRNP H2 (magenta), Homer1 (cyan) and MAP2 (grey). Merged images are shown. **(G–J)** Quantification of the percentage of postsynaptic (Homer1-positive) puncta containing hnRNP H2 (G), postsynaptic (Homer1-positive) hnRNP H2 puncta density (H), integrated density per dendritic length (I), and puncta area per dendritic length (J). Data are expressed relative to control and presented as mean ± SEM from 3 independent neuronal preparations, for western blot analysis, and 4 independent neuronal preparations for immunocytochemistry analysis. At least 12 fields containing multiple dendrites were analysed per condition in each experiment. Statistical significance was determined using two-tailed unpaired Student’s *t*-tests. p < 0.05 (*), p < 0.01(**), p < 0.001(***), p < 0.0001(****). Scale bars: (B) 5 μm; inset, 2 μm; (C) 2 μm; (D) 10 μm; inset, 0.5 μm; (F) 2.5 μm.

To determine whether synaptic activity modulates the dendritic distribution of hnRNP H2, we stimulated cultured hippocampal neurons using a well-established chemical protocol consisting of bicuculline (Bic), 4-aminopyridine (4-AP), and glycine (Gly) (M. V. Caldeira et al., 2013) to enhance excitatory synaptic transmission **(Fig. 1E, F)**. Quantitative analysis revealed that neuronal activity significantly increased the proportion of Homer1-positive sites colocalising with hnRNP H2 **(Fig. 1G)**. Moreover, the number, integrated density (IntDen), and area of postsynaptic hnRNP H2 puncta were all markedly elevated following stimulation **(Fig. 1H-J)**. Additionally, we observed a significant increase in dendritic hnRNP H2 levels **(Fig. S1B-D)**, further supporting activity-dependent redistribution of the protein. Notably, these changes occurred without alterations in Homer1 levels **(Fig. S1E-G)**, indicating a specific upregulation of hnRNP H2 at synaptic sites.

Together, these data demonstrate that hnRNP H2 is expressed in hippocampal neurons during development and localises to the nucleus, cytoplasm, and dendrites. Importantly, we show that hnRNP H2 is also present at excitatory synapses and undergoes an activity-dependent recruitment to these postsynaptic compartments.

### hnRNP H2 is required for dendritic development and cortical neuron migration

To investigate the role of hnRNP H2 in hippocampal neuronal development, we generated a lentiviral plasmid co-expressing GFP together with a miR30-based short hairpin RNA (shRNA) targeting endogenous rodent *Hnrnph2* mRNA. A matched GFP-expressing vector lacking the shRNA insert was used as a control (Ctr). Transfection of cultured rat hippocampal neurons with the hnRNP H2 knockdown (KD) construct led to a significant reduction in dendritic hnRNP H2 levels at both DIV 7 and DIV 14 **(Fig. S2A-J)**, confirming efficient knockdown at early and later developmental stages. To assess specificity and rule out off-target effects, we immunostained for hnRNP H1, a closely related functional paralog. Notably, hnRNP H2 knockdown had no effect on dendritic hnRNP H1 levels **(Fig. S2K-O)**, validating the specificity of the genetic silencing approach.

Having established the efficacy and specificity of the knockdown system, we next employed it to investigate the role of hnRNP H2 in regulating hippocampal neuronal morphology **(Fig. 2A, H)**. At DIV 7, hnRNP H2 knockdown significantly reduced total dendritic length, without affecting the number of primary dendrites, branch points, or terminal branches **(Fig. 2B-G)**. In contrast, by DIV 14, knockdown of hnRNP H2 led to increased dendritic complexity **(Fig. 2I-J)**. At this later developmental stage, total dendritic length and the number of branch points remained unchanged; however, the number of primary and terminal dendritic branches was significantly elevated compared to control neurons **(Fig. 2K-N)**. Importantly, hnRNP H2 depletion also resulted in a significant enlargement of the soma at DIV 14, an effect not observed at DIV 7 **(Fig. S3)**. We next assessed whether hnRNP H2 influences dendritic spine development **(Fig. 2O-Q)**. Although total spine density was unchanged, hnRNP H2 knockdown significantly reduced the density and proportion of mushroom spines, typically indicative of mature synapses. While changes in other spine subtypes did not reach statistical significance, there was a trend towards an increased density and proportion of filopodia, together with trends towards reduced densities and fraction of branched, stubby and thin spines **(Fig. 2R, S)**. Consistent with this, classification of spines into mature and immature categories revealed a selective reduction in the density of mature spines following hnRNP H2 knockdown **(Fig. 2T).**

**Figure 2.**
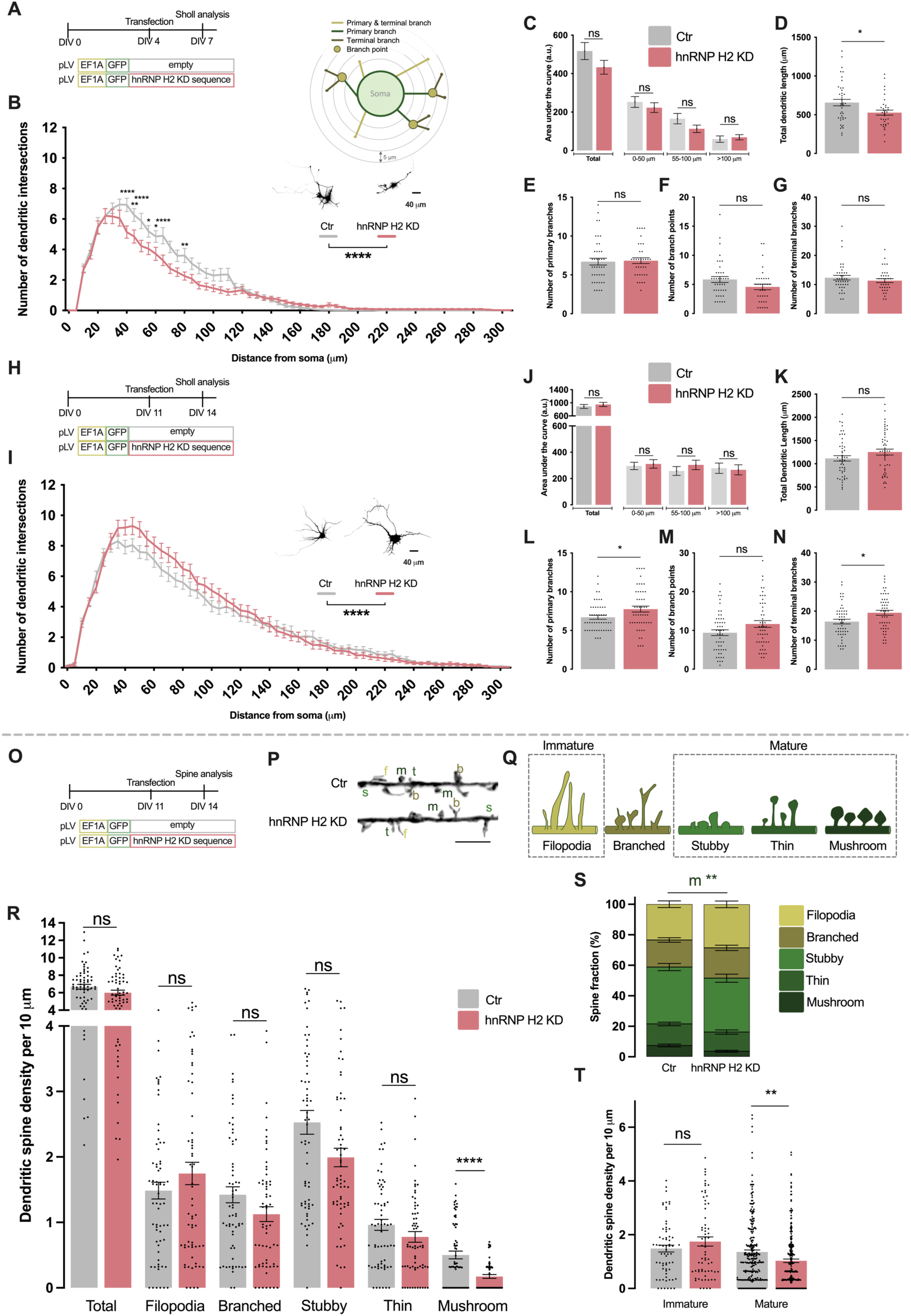
hnRNP H2 knockdown induces structural and dendritic spine alterations in developing and mature hippocampal neurons. **(A)** Experimental paradigm for the analysis of neuronal morphology in developing hippocampal neurons. Primary rat hippocampal neurons were transfected at DIV 4 with either a control plasmid (Ctr) or a plasmid expressing an hnRNP H2-targeting knockdown sequence (hnRNP H2 KD sequence). Neurons were fixed at DIV 7 and immunostained for MAP2 to visualize dendritic arbors, while GFP identified transfected neurons. Schematic representation of the parameters evaluated by Sholl analysis. **(B–G)** Quantification of neuronal complexity in developing hippocampal neurons at DIV 7, including Sholl analysis showing the number of dendritic intersections as a function of distance from the soma (B), area under the Sholl curve (C), total dendritic length (D), number of primary branches (E), number of branch points (F), and number of terminal branches (G). **(H)** Experimental paradigm for the analysis of neuronal morphology in mature hippocampal neurons. Primary rat hippocampal neurons were transfected at DIV 11 with either the Ctr or hnRNP H2 KD sequence plasmid and fixed at DIV14 for MAP2 immunostaining. **(I–N)** Quantification of neuronal complexity in mature hippocampal neurons at DIV 14, including Sholl analysis (I), area under the Sholl curve (J), total dendritic length (K), number of primary branches (L), number of branch points (M), and number of terminal branches (N). **(O)** Experimental paradigm for dendritic spine analysis. Hippocampal neurons were transfected at DIV 11 with either the Ctr or hnRNP H2 KD sequence plasmid and fixed at DIV 14. Cultures were immunostained for GFP to enhance the signal and visualize dendritic spines. **(P)** Representative images of dendritic segments from Ctr and hnRNP H2-knockdown neurons displaying multiple dendritic spines. f – filopodia, b – branched, s – stubby, t – thin, m – mushroom. **(Q)** Schematic representation of the dendritic spine categories used for morphological classification (filopodia, branched, stubby, thin and mushroom). Branched spines were considered a transitional morphology and were excluded from the immature and mature spine groups. **(R–T)** Quantification of dendritic spine density per dendritic length and distribution of dendritic spine subtypes (R), dendritic spine density normalized to fractional counts (S), and dendritic spine density according to maturation stage (T). Spine maturation was assessed by classifying spines as immature (filopodia) or mature (thin, stubby and mushroom). Data are expressed relative to control and presented as mean ± SEM from 3–4 independent neuronal preparations. At least 12 neurons per condition, in each experiment, were analysed for neuronal morphology, and a total of 65 dendrites per condition were analysed for spine morphology. Statistical significance was determined using two-way ANOVA followed by Sidak’s multiple comparisons test for Sholl analysis. For all other comparisons, data distribution was first assessed for normality. Normally distributed datasets were analysed using two-tailed unpaired Student’s *t*-tests, whereas non-normally distributed datasets were analysed using two-tailed Mann–Whitney tests. p < 0.05 (*), p < 0.01 (**), p < 0.0001 (****); ns, not statistically significant. Scale bar: (B, I) 40 μm; (P) 2.5 μm.

To extend these findings *in vivo*, we assessed the effects of hnRNP H2 depletion on cortical neuron migration during embryonic development. *In utero* electroporation (IUE) was performed at embryonic day 15.5 (E15.5) to introduce either the control or hnRNP H2 knockdown constructs into the developing mouse cortex **(Fig. 3A-B)**. Analysis at E18 revealed a significant migration defect in hnRNP H2-depleted neurons, with a reduced proportion of GFP-positive cells reaching the cortical plate (CP) and intermediate zone (IZ) **(Fig. 3B, C)** and a corresponding accumulation in the ventricular/subventricular zones (VZ/SVZ) **(Fig. 3B, D).**

**Figure 3.**
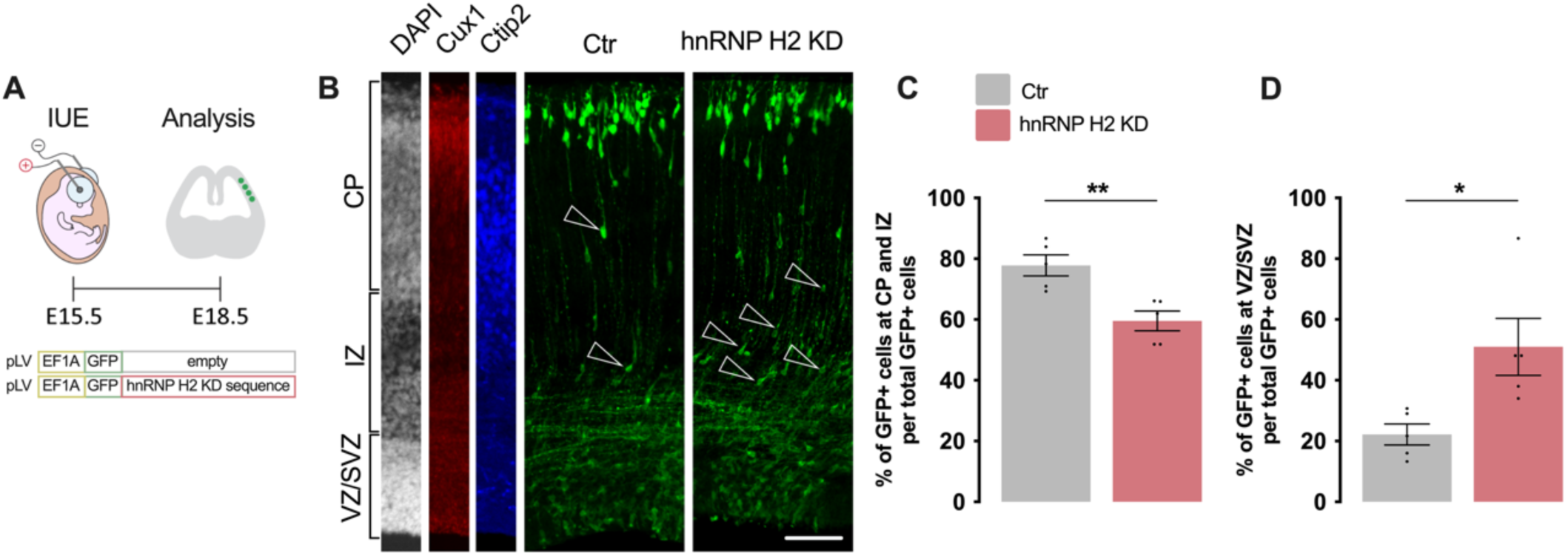
hnRNP H2 knockdown impairs neuronal migration during corticogenesis *in vivo*. **(A)** Experimental paradigm for the analysis of neuronal migration by in utero electroporation (IUE). Mouse embryos were *in utero* electroporated at embryonic day 15.5 (E15.5) with either a control plasmid (Ctr) or a plasmid expressing an hnRNP H2-targeting knockdown sequence (hnRNP H2 KD sequence). Embryos were collected at E18, and coronal brain sections were immunostained for Ctip2 and Cux1 and counterstained with DAPI. GFP fluorescence was used to identify electroporated neurons. The schematic illustrates the IUE procedure targeting progenitors in the embryonic somatosensory cortex. **(B)** Representative confocal images of coronal sections from E18 mouse cortex electroporated with Ctr or hnRNP H2 KD sequence plasmids. GFP-positive neurons (green) are shown together with DAPI (grey), and Cux1 (red) and Ctip2 (blue). White arrowheads indicate ectopic GFP-positive neurons in the intermediate zone (IZ) and lower cortical plate (CP). Ventricular/subventricular zone (VZ/SVZ) is indicated. **(C)** Quantification of the percentage of GFP-positive neurons located within CP and IZ relative to the total number of GFP-positive cells in the same coronal section. **(D)** Quantification of the percentage of GFP-positive neurons located within VZ/SVZ relative to the total number of GFP-positive cells in the same coronal section. Data are expressed relative to control and presented as mean ± SEM from 3 independent experiments, with at least one brain section analysed per animal. Statistical significance was determined using a two-tailed unpaired Student’s *t*-test. p < 0.05 (*), p < 0.01 (**). Scale bar: (B) 200 μm.

Together, these results identify hnRNP H2 as a key regulator of neuronal development, with critical roles in shaping dendritic architecture, promoting dendritic spine maturation, and ensuring proper cortical neuron migration, all of which are critical for normal brain development.

### hnRNP H2 regulates excitatory synaptic transmission

Given the results of Fig. 1 showing that hnRNP H2 localises to the synaptic compartment, and in light of established roles for hnRNP family proteins in regulating excitatory synaptic function (Afonso et al., 2019; Jung et al., 2020; Leal et al., 2017), we next investigated whether hnRNP H2 modulates postsynaptic AMPA receptor expression. We assessed the surface and synaptic distribution of the GluA1 AMPA receptor subunit using immunocytochemistry with an antibody targeting its extracellular domain **(Fig. 4A, B)** and colocalisation with the postsynaptic marker PSD-95. Labelling of the surface receptors was performed under non-permeabilizing conditions. While hnRNP H2 knockdown did not alter the percentage of GluA1-positive PSD-95 puncta **(Fig. 4C)**, the number, integrated density (IntDen), and area of GluA1 puncta at postsynaptic sites were significantly reduced **(Fig. 4D-F)**. In addition, total surface levels of GluA1 were markedly decreased following hnRNP H2 depletion **(Fig. S4)**, indicating reduced GluA1-containing AMPARs at both synaptic and extrasynaptic sites. These effects occurred without detectable changes in overall PSD-95 levels **(Fig. 4G-J)**, suggesting a selective role for hnRNP H2 in regulating surface GluA1 expression.

**Figure 4.**
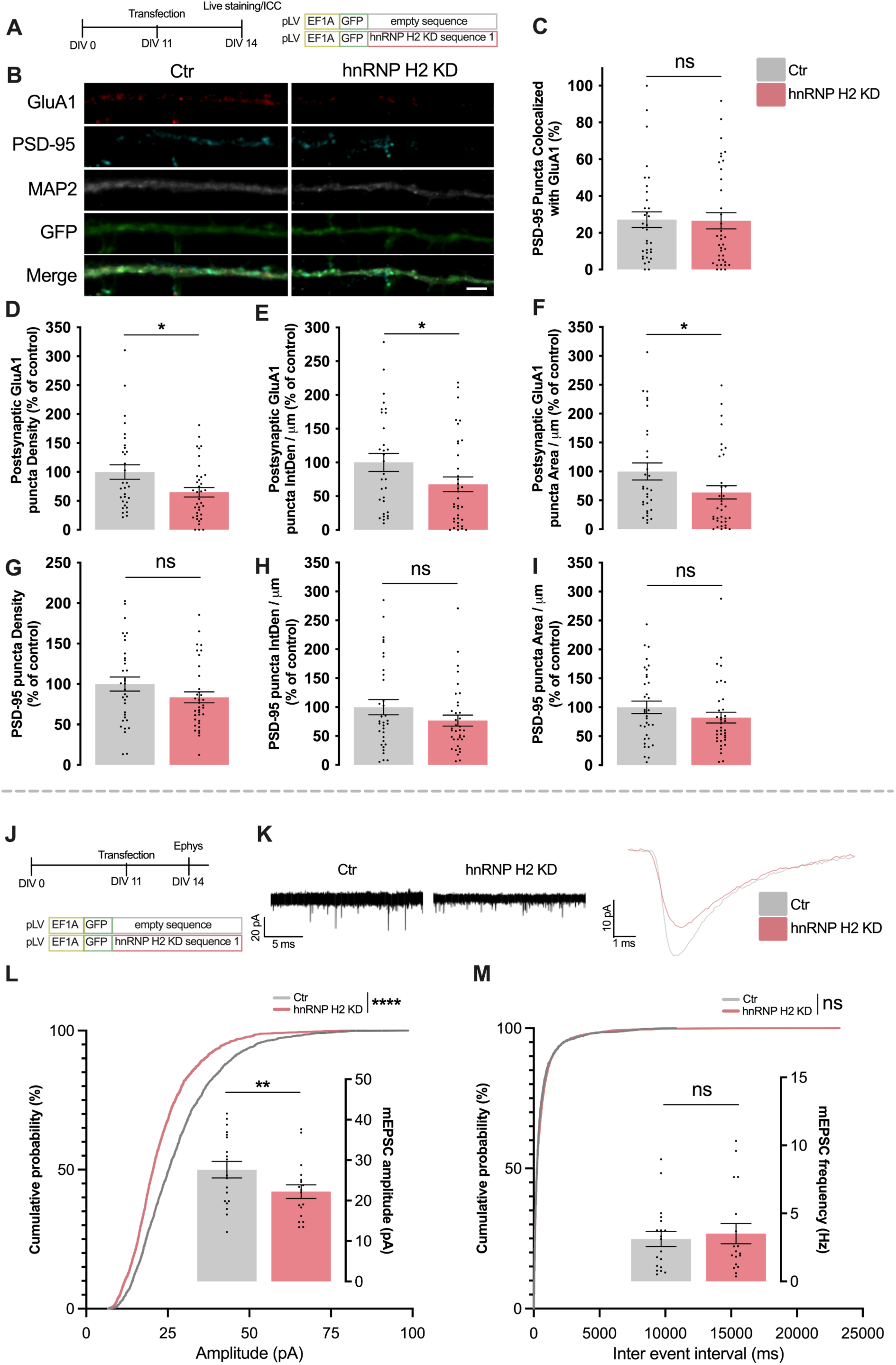
hnRNP H2 knockdown reduces AMPA receptor surface expression and impairs basal excitatory synaptic transmission in cultured hippocampal neurons. **(A)** Experimental paradigm for the analysis of AMPA receptor (AMPAR) surface expression. Primary rat hippocampal neurons were transfected at DIV 11 with either a control plasmid (Ctr) or a plasmid expressing an hnRNP H2-targeting knockdown sequence (hnRNP H2 KD sequence). Neurons were fixed at DIV 14 and immunostained for the AMPAR subunit GluA1, PSD-95 and MAP2. GFP was used to identify transfected neurons. **(B)** Representative images of dendritic segments from Ctr and hnRNP H2-knockdown neurons immunostained for surface GluA1 (red), PSD-95 (cyan) and MAP2 (grey). Merged images are shown. **(C–F)** Quantification of the percentage of postsynaptic (PSD-95-positive) puncta containing surface GluA1 (C), postsynaptic (PSD-95-positive) surface GluA1 puncta density (D), integrated density per dendritic length (E), and puncta area per dendritic length (F). **(G–I)** Quantification of PSD-95 puncta density (G), integrated density per dendritic length (H), and puncta area per dendritic length (I). **(J)** Experimental paradigm for electrophysiological recordings of AMPAR-mediated miniature excitatory postsynaptic currents (mEPSCs). Rat hippocampal neurons were transfected at DIV 11 with Ctr or a hnRNP H2 KD sequence, and whole-cell recordings were performed at DIV 14. AMPAR-mediated mEPSCs were pharmacologically isolated using tetrodotoxin (TTX), bicuculline (Bic) and D-AP5. GFP fluorescence was used to identify transfected neurons. **(K)** Representative traces of AMPAR-mediated mEPSCs (left) and averaged single mEPSC events (right) recorded from Ctr and hnRNP H2 KD neurons. **(L, M)** Quantification of the cumulative probability distributions and mean values of mEPSC amplitude (L) and frequency (M). Data are expressed relative to control and presented as mean ± SEM. AMPAR surface expression analyses represent 3 independent experiments from different neuronal preparations. Electrophysiological recordings were obtained from at least 5 independent neuronal preparations. Statistical significance was determined using two-tailed Mann–Whitney tests (mean frequency), two-tailed unpaired Student’s *t*-tests (mean amplitude), and Kolmogorov–Smirnov tests for cumulative probability distributions. p < 0.05 (*), p < 0.01 (**), p < 0.0001 (****); ns, not statistically significant. Scale bar: (B) 2.5 μm.

To determine whether these molecular changes impact excitatory synaptic transmission, we next performed whole-cell patch-clamp recordings to measure AMPA receptor-mediated miniature excitatory postsynaptic currents (mEPSCs) in transfected cultured hippocampal neurons **(Fig. 4J, K)**. Given the low rate of transfection using the Ca^2+^-phosphate method, the observed alterations in the mEPSCs result from the downregulation of hnRNP H2 in the postsynaptic neuron that is recorded (Leal et al., 2017). hnRNP H2 knockdown led to a significant reduction in mEPSC amplitude **(Fig. 4L)**, consistent with a decrease in postsynaptic AMPA receptor content. In contrast, mEPSC frequency remained unchanged **(Fig. 4M)**, indicating that presynaptic release probability and synapse number were not substantially affected. This selective reduction in mEPSC amplitude aligns with the immunocytochemistry data showing decreased GluA1 levels at postsynaptic sites.

Together, these findings identify hnRNP H2 as a key regulator of synaptic GluA1-containing AMPA receptors, revealing a previously unrecognized role for this RBP in maintaining excitatory synaptic strength in hippocampal neurons.

### Pathogenic hnRNP H2 variants affect the neuronal localisation of hnRNP H2

*HNRNPH2*-related neurodevelopmental disorder is caused by pathogenic variants that predominantly cluster within the qRRM and NLS domains of the hnRNP H2 protein (Gillentine et al., 2021; Bain et al., 2016, 2021; Somashekar et al., 2020; White-Brown et al., 2022; Harmsen et al., 2019; Jepsen et al., 2019; Peron et al., 2020; Kreienkamp et al., 2022; Lourinho et al., 2025) **(Fig. 5A)**. To investigate how these disease-associated variants affect hnRNP H2 structure *in silico*, we first performed molecular dynamics (MD) simulations over a 500-nanosecond (ns) trajectory on the wild-type (WT) protein and on disease-associated variants affecting either the NLS (p.R206W, p.R206G, and p.R206Q) or qRRM (p.R114W) domains. Conformational dynamics were assessed by tracking the distance between the two qRRM domains over time, providing a measure of protein flexibility and compactness. Among the NLS-domain variants, p.R206G showed the most stable behaviour, rapidly adopting a compact conformation that remained consistent throughout the simulation **(Fig. 5B)**. The p.R206W variant also led to a more compact and less flexible structure compared to WT, although to a lesser extent than p.R206G **(Fig. 5B)**. In contrast, the p.R206Q variant exhibited a broader range of motion and greater conformational variability, indicating increased flexibility **(Fig. 5B)**. Interestingly, the qRRM-domain variant p.R114W also resulted in enhanced conformational stability relative to the WT protein **(Fig. 5B)**. Collectively, these simulations demonstrate that pathogenic variants differentially alter the conformational dynamics of hnRNP H2, potentially impacting its functional properties in neurons.

**Figure 5.**
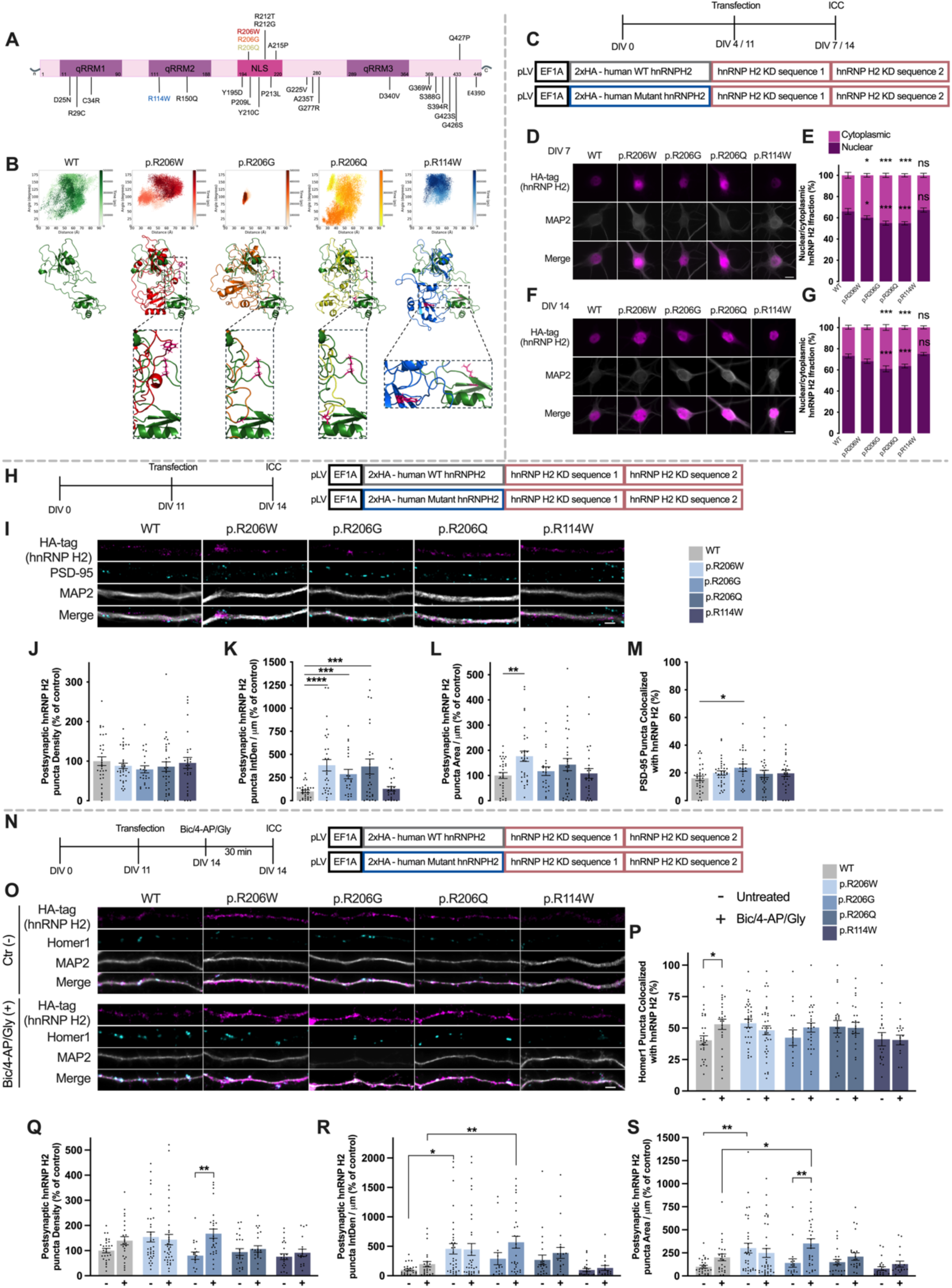
Disease-associated hnRNP H2 variants disrupt hnRNP H2 conformation, subcellular localisation and activity-dependent synaptic recruitment. **(A)** Schematic representation of the HNRNPH2 gene showing the location of pathogenic variants associated with HNRNPH2-related neurodevelopmental disorder. **(B)** Molecular dynamics simulations of human hnRNP H2 WT or disease-associated hnRNP H2 variants (p.R114W, p.R206Q, p.R206W or p.R206G). Upper panels, conformational landscapes showing the trajectories sampled during 500 ns simulations, represented as the distance between qRRM1 and qRRM2 plotted against the interdomain angle. Lower panels, representative structural models of the most populated conformations aligned to the WT, illustrating mutation-induced changes in the orientation of the qRRM2 and NLS domains. **(C)** Experimental paradigm for the analysis of hnRNP H2 nucleocytoplasmic localisation in developing hippocampal neurons. Primary rat hippocampal neurons were transfected at DIV 4 or DIV 7 with bicistronic lentiviral plasmids encoding HA-tagged human hnRNP H2 wild-type (WT) or the disease-associated variants p.R114W, p.R206Q, p.R206W or p.R206G, together with two knockdown sequences targeting endogenous rodent *Hnrnph2* mRNA. Neurons were fixed at DIV 7 or DIV 14 and immunostained for HA and MAP2. **(D–G)** Representative images of neuronal cell bodies immunostained for HA-tagged hnRNP H2 (magenta) and MAP2 (grey), and quantification of hnRNP H2 subcellular localisation in developing (DIV 7; D, E) and mature (DIV 14; F, G) neurons. hnRNP H2 fluorescence intensity was quantified separately in the nuclear and cytoplasmic compartments of the neuronal soma (E, G), and the relative nuclear and cytoplasmic fractions were calculated based on the total hnRNP H2 fluorescence signal measured across both compartments. **(H)** Experimental paradigm for the analysis of dendritic and synaptic hnRNP H2 localisation. Hippocampal neurons were transfected at DIV 11 with bicistronic lentiviral plasmids encoding HA-tagged human hnRNP H2 wild-type (WT) or the disease-associated variants p.R114W, p.R206Q, p.R206W or p.R206G, together with two knockdown sequences targeting endogenous rodent hnRNP H2, fixed at DIV 14 and immunostained for HA-tagged hnRNP H2, PSD-95 and MAP2. **(I)** Representative images of dendritic segments from neurons expressing hnRNP H2 human WT or hnRNP H2 variants immunostained for HA-tagged hnRNP H2 (magenta), PSD-95 (cyan) and MAP2 (grey). Merged images are shown. **(J–M)** Quantification of postsynaptic hnRNP H2 puncta density (J), integrated density per dendritic length (K) and puncta area per dendritic length (L). Synaptic localisation was further assessed by quantifying the percentage of PSD-95-positive puncta containing hnRNP H2 (M). **(N)** Experimental paradigm for activity-dependent synaptic recruitment. Hippocampal neurons expressing WT or mutant hnRNP H2 were stimulated at DIV 14 with bicuculline (50 μM), 4-aminopyridine (2.5 mM) and glycine (10 μM) for 30 min before fixation and immunostaining for HA, Homer1 and MAP2. **(O)** Representative images of dendritic segments from neurons from unstimulated and stimulated neurons expressing hnRNP H2 human WT or hnRNP H2 variants immunostained for HA-tagged hnRNP H2 (magenta), Homer1 (cyan) and MAP2 (grey). Merged images are shown. (**P–S**) Quantification of the percentage of Homer1-positive puncta containing hnRNP H2 (P), postsynaptic (Homer1-positive) hnRNP H2 puncta density (Q), integrated density per dendritic length (R) and puncta area per dendritic length (S). Data are expressed relative to control and presented as mean ± SEM from 3 independent neuronal preparations for localisation analyses and 4 independent neuronal preparations for activity-dependent recruitment experiments. At least 12 neurons or 12 fields of view containing multiple dendrites were analysed per condition in each independent experiment. Statistical significance was determined using two-tailed unpaired Student’s *t*-tests for localisation analyses and two-way ANOVA followed by Tukey’s multiple comparisons test for activity-dependent recruitment. p < 0.05 (*), p < 0.01 (**), p < 0.001 (***), p < 0.0001 (****); ns, not statistically significant. Scale bars: (D, F) 10 μm; (I, O) 2.5 μm.

To investigate the neuronal consequences of these variants in cultured hippocampal neurons, we developed a molecular replacement strategy using a single bicistronic lentiviral plasmid to simultaneously knock down endogenous rodent *Hnrnph2* mRNA and express HA-tagged human hnRNP H2 (WT or mutant) in its place. To validate that this approach preserves physiological levels and localisation of hnRNP H2, we compared dendritic hnRNP H2 abundance in neurons transfected with the WT construct to that of mock- and non-transfected controls **(Fig. S5A, B).** HA-tagged human hnRNP H2 was selectively expressed in transfected cells and exhibited predominantly nuclear distribution, consistent with endogenous hnRNP H2. Quantitative fluorescence analysis revealed no significant differences in the number, integrated density, or area of dendritic hnRNP H2 puncta across conditions **(Fig. S5C-E)**. These results confirm that our molecular replacement strategy maintains hnRNP H2 expression within physiological parameters, validating it, along with all variant constructs, for subsequent functional investigations.

The subcellular distribution of hnRNP H2 is governed by tightly regulated nucleocytoplasmic transport, mediated by the interaction between its NLS and the import receptor Karyopherin-β2 (Kapβ2) (Gonzalez et al., 2023). Having confirmed that our molecular replacement system supports physiologically faithful expression of both WT and mutant HA-tagged human hnRNP H2, we next assessed how disease-associated variants affect its localisation in developing hippocampal neurons **(Fig. 5C-G)**. At DIV 7, human hnRNP H2 fluorescence intensity was quantified separately in the nuclear and cytoplasmic compartments of the neuronal soma, and the relative nuclear and cytoplasmic fractions were calculated based on the total hnRNP H2 fluorescence signal measured across these two compartments. WT hnRNP H2 localised predominantly to the nucleus (∼66%), with a smaller cytoplasmic fraction (∼34%) **(Fig. 5D, E)**. In contrast, neurons expressing any of the NLS-domain variants (p.R206W, p.R206G, or p.R206Q), but not the qRRM-domain variant p.R114W, exhibited a marked redistribution of hnRNP H2 from the nucleus to the cytoplasm **(Fig. 5D, E, Fig. S6A-D)**. Morphological analysis at DIV 7 further showed that neurons expressing p.R206W or p.R114W hnRNP H2 exhibited significantly increased nuclear area relative to WT, while soma size was selectively increased in p.R114W-expressing neurons **(Fig. S6E, F)**. At DIV 14, analysis of hnRNP H2 fluorescence intensity in the nuclear and cytoplasmic compartments similarly revealed a predominantly nuclear distribution of WT hnRNP H2, with ∼73% of the total nuclear plus cytoplasmic signal detected in the nucleus and ∼27% in the cytoplasm **(Fig. 5F, G)**. Consistent with earlier observations, NLS mutants displayed persistently reduced nuclear localisation and increased cytoplasmic levels, while the distribution of the p.R114W variant remained indistinguishable from WT **(Fig. 5F, G, Fig. S6G-J)**. At this later stage, no significant changes in nuclear or soma size were observed across any condition **(Fig. S6K, L)**.

Given our earlier observation that endogenous hnRNP H2 localises to dendrites and synapses, we next asked whether the altered nucleocytoplasmic distribution observed in NLS mutants also affected its localisation within these compartments **(Fig. 5H, I)**. Despite a reduction in dendritic hnRNP H2 puncta number, all NLS-domain variants exhibited increased integrated intensity, consistent with protein accumulation within dendrites rather than uniform distribution **(Fig. S6M, N)**. This effect was absent in neurons expressing the p.R114W variant, in agreement with its normal nucleocytoplasmic distribution **(Fig. S6M, N)**. Among the NLS mutants, p.R206W induced the most pronounced accumulation, as reflected by a significant increase in dendritic puncta area **(Fig. S6O)**. Analysis of postsynaptic hnRNP H2 puncta revealed no differences in puncta number between neurons expressing WT or mutant variants **(Fig. 5J)**. However, all NLS mutants showed a significant increase in postsynaptic hnRNP H2 integrated intensity, indicative of protein accumulation at synaptic sites, an effect completely absent in neurons expressing the p.R114W variant **(Fig. 5K)**. Notably, p.R206W again exhibited the strongest phenotype, with a significant increase in postsynaptic hnRNP H2 puncta area relative to WT **(Fig. 5L)**. Furthermore, the percentage of PSD-95-positive sites containing hnRNP H2 was elevated in NLS mutants, reaching statistical significance for the p.R206G variant **(Fig. 5M).** Notably, PSD-95 puncta exhibited only minor alterations across conditions, with reduced density and area observed in p.R206G-expressing neurons **(Fig. S6P-R).**

Our initial findings demonstrated that synaptic stimulation enhances dendritic and postsynaptic localisation of endogenous hnRNP H2. Building on these observations, we next assessed whether disease-associated variants in hnRNP H2 disrupt its activity-dependent redistribution **(Fig. 5N, O)**. Under basal conditions, none of the hnRNP H2 variants significantly altered dendritic puncta number relative to WT **(Fig. S6S)**. However, neurons expressing NLS mutants exhibited elevated integrated hnRNP H2 intensity, with p.R206W additionally showing increased puncta area **(Fig. S6T, U)**, consistent with aberrant accumulation of the protein in dendritic compartments. These alterations were not observed for the p.R114W variant, in line with the results described above **(Fig. S6S-U)**. Following stimulation with Bic/4-AP/Gly, WT hnRNP H2 displayed robust activity-dependent increases in dendritic puncta number and area **(Fig. S6S-U)**, mirroring the behavior of endogenous hnRNP H2 **(Fig. S1B-D)**. At the postsynaptic level, synaptic stimulation increased the proportion of Homer1-positive sites colocalising with WT hnRNP H2 **(Fig. 5P)**, alongside a trend toward higher number, integrated intensity, and area of postsynaptic hnRNP H2 puncta, supporting the activity-dependent recruitment of WT hnRNP H2 to synaptic compartments **(Fig. 5Q-S)**. The lack of statistical significance is likely due to detection being limited to exogenous HA-tagged human hnRNP H2, excluding the endogenous protein pool also recruited to synapses. In contrast to WT-expressing neurons, the p.R206W variant failed to show any increase in dendritic or postsynaptic hnRNP H2 following stimulation. Indeed, neither the fraction of Homer1-positive puncta colocalising with hnRNP H2 nor the postsynaptic hnRNP H2 parameters changed with treatment **(Fig. 5Q-S, Fig. S6S-U)**, suggesting that the p.R206W variant drives a constitutive mislocalisation of hnRNP H2 that occludes further recruitment. p.R206G-expressing neurons, by contrast, retained partial responsiveness, with stimulation inducing significant increases in both total dendritic and postsynaptic hnRNP H2 **(Fig. 5Q-S, Fig. S6S-U).** The **p.R206**Q variant showed a similar intermediate phenotype, with modest yet detectable dendritic and postsynaptic increases of hnRNP H2 upon stimulation **(Fig. 5Q-S, Fig. S6S-U)**. Notably, the p.R114W variant, despite lacking basal dendritic or synaptic accumulation, also failed to respond to stimulation, with hnRNP H2 distribution remaining unchanged between basal and treated conditions **(Fig. 5Q–S, Fig. S6S-U)**. By contrast, Homer1 puncta levels remained largely unchanged across all genotypes **(Fig. S6V-Z)**, suggesting that activity primarily alters postsynaptic hnRNP H2 localisation without detectably affecting Homer1-positive postsynaptic sites

Overall, our findings show that pathogenic variants in the NLS of hnRNP H2 impair its nuclear import, causing abnormal cytoplasmic and dendritic accumulation in hippocampal neurons. This mislocalisation extends to synapses, disrupting the normal activity-dependent recruitment of hnRNP H2. Interestingly, the p.R114W variant similarly impairs this dynamic synaptic targeting of hnRNP H2 despite not affecting the nuclear localisation sequence of the protein.

### hnRNP H2 variants impair dendritic architecture and spine development

Altered dendritic architecture is a hallmark of numerous neurodevelopmental and neuropsychiatric disorders, as evidenced by both human studies and animal models (G. L. Caldeira et al., 2022; Edfawy et al., 2019; Forrest et al., 2018). Building on our findings that hnRNP H2 is essential for dendritic morphogenesis and spine maturation, we next examined how disease-associated variants in *HNRNPH2* affect these processes across key stages of hippocampal neuronal development *in vitro* **(Fig. 6A, H)**.

**Figure 6.**
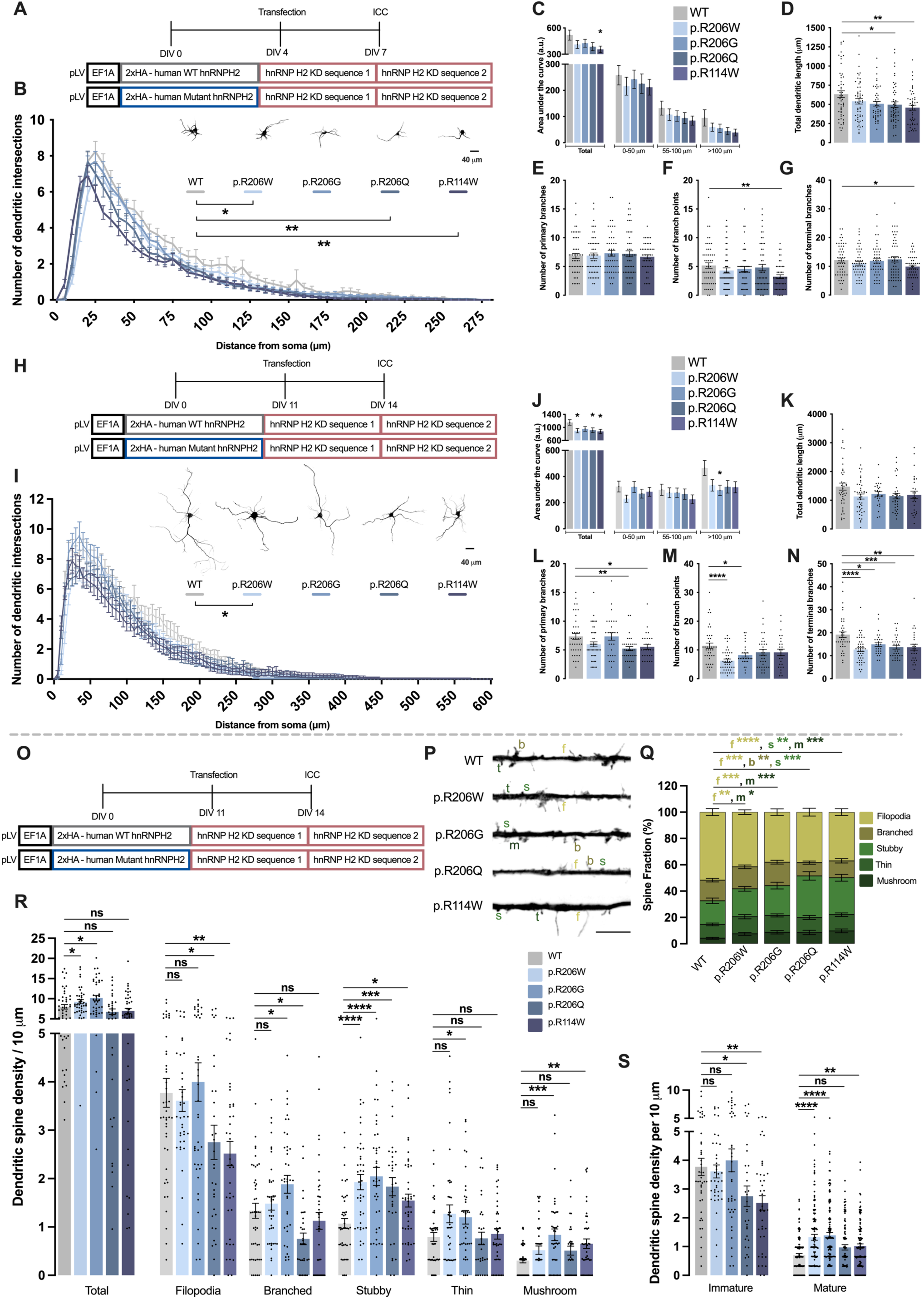
Disease-associated hnRNP H2 variants impair dendritic development and alter dendritic spine maturation in hippocampal neurons. **(A)** Experimental paradigm for the analysis of neuronal morphology in developing hippocampal neurons. Primary rat hippocampal neurons were transfected at DIV 4 with bicistronic lentiviral plasmids expressing HA-tagged human modified hnRNP H2 wild-type (WT) or disease-associated hnRNP H2 variants (p.R114W, p.R206Q, p.R206W or p.R206G) together with two knockdown sequences targeting endogenous rodent hnRNP H2. Neurons were fixed at DIV 7 and immunostained for MAP2 to visualize dendritic arbors, while the HA tag identified transfected neurons. Schematic representation of the parameters evaluated by Sholl analysis. **(B–G)** Quantification of neuronal complexity in developing hippocampal neurons at DIV 7, including Sholl analysis showing the number of dendritic intersections as a function of distance from the soma (B), area under the Sholl curve (C), total dendritic length (D), number of primary branches (E), number of branch points (F), and number of terminal branches (G). **(H)** Experimental paradigm for the analysis of neuronal morphology in mature hippocampal neurons. Primary rat hippocampal neurons were transfected at DIV 11 with bicistronic lentiviral plasmids encoding HA-tagged human hnRNP H2 wild-type (WT) or the disease-associated variants p.R114W, p.R206Q, p.R206W or p.R206G, together with two knockdown sequences targeting endogenous rodent hnRNP H2. Neurons were fixed at DIV 14 for MAP2 immunostaining. The HA tag was used to identify transfected neurons. **(I–N)** Quantification of neuronal complexity in mature hippocampal neurons at DIV 14, including Sholl analysis (I), area under the Sholl curve (J), total dendritic length (K), number of primary branches (L), number of branch points (M), and number of terminal branches (N). **(O)** Experimental paradigm for dendritic spine analysis. Hippocampal neurons were transfected at DIV 11 with bicistronic lentiviral plasmids encoding HA-tagged human hnRNP H2 wild-type (WT) or the disease-associated variants p.R114W, p.R206Q, p.R206W or p.R206G, together with two knockdown sequences targeting endogenous rodent hnRNP H2. A separate GFP-expressing plasmid was co-transfected to enhance neuronal morphology visualization. Neurons were fixed at DIV 14 and immunostained for GFP to enhance the signal. **(P)** Representative images of dendritic segments from WT and mutant hnRNP H2-expressing hippocampal neurons. f, filopodia; b, branched; s, stubby; t, thin; m, mushroom. **(Q–S)** Quantification of dendritic spine density normalized to fractional counts (Q), dendritic spine density per dendritic length and distribution of dendritic spine subtypes (R), and dendritic spine density according to maturation stage (S). Spine maturation was assessed by classifying spines as immature (filopodia) or mature (thin, stubby and mushroom). Data are expressed relative to control and presented as mean ± SEM from 3–5 independent neuronal preparations. At least 12–15 neurons per condition, in each experiment, were analysed for neuronal morphology. For dendritic spine analyses, at least 10 dendrites from different neurons per condition were analysed in each experiment. Statistical significance was determined using two-way ANOVA followed by Sidak’s multiple comparisons test for Sholl analysis. For all other comparisons, data distribution was first assessed for normality. Normally distributed datasets were analysed using two-tailed unpaired Student’s *t*-tests, whereas non-normally distributed datasets were analysed using two-tailed Mann– Whitney tests. p < 0.05 (*), p < 0.01 (**), p < 0.001 (***), p < 0.0001 (****); ns, not statistically significant. Scale bars: (B, I) 40 μm; (P) 5 μm.

At DIV 7, all hnRNP H2 variants, except for p.R206G, significantly reduced overall neuronal complexity relative to WT-expressing neurons **(Fig. 6B)**, as assessed by Sholl analysis, with the most pronounced effect observed for the p.R114W variant. This was reflected by a statistically significant decrease in the total area under the curve (AUC) compared with WT-expressing neurons **(Fig. 6C)**. Although all variants showed a trend toward reduced total dendritic length, only the p.R206Q and p.R114W variants produced statistically significant effects **(Fig. 6D)**. Notably, p.R114W-expressing neurons also exhibited a significant decrease in the number of branch points and terminal branches **(Fig. 6E-G)**. By DIV 14, the impact of hnRNP H2 variants on dendritic architecture was more heterogeneous. Only the p.R206W variant significantly reduced overall neuronal complexity compared to WT **(Fig. 6I)**, while the p.R206W, p.R206Q and p.R114W variants displayed a significantly reduced total AUC **(Fig. 6J)**. While total dendritic length remained unchanged across all conditions at this stage, neurons expressing p.R206Q and p.R114W variants showed a reduced number of primary dendrites, and both the p.R206W and p.R206G variants significantly decreased branch point number **(Fig. 6K-M)**. Importantly, all neurons expressing hnRNP H2 mutants exhibited a significant reduction in terminal branch number compared to WT-expressing cells **(Fig. 6N)**.

We next investigated the impact of hnRNP H2 variants on dendritic spine development **(Fig. 6O)**. Morphological analysis revealed distinct, variant-specific alterations in spine architecture and density. All hnRNP H2 variants resulted in a significant reduction in the proportion of immature, filopodia-like spines, accompanied by a corresponding increase in mature spine subtypes, most notably stubby and mushroom spines **(Fig. 6P, Q)**. Consistently, filopodia spine density was markedly decreased in neurons expressing the p.R206Q and p.R114W variants, whereas the density of stubby, thin, and mushroom spines was generally elevated across all mutant conditions **(Fig. 6R)**. Notably, total spine density was significantly increased only in neurons expressing the p.R206W and p.R206Q variants **(Fig. 6R)**. Analysis of immature and mature spines revealed a selective reduction in immature spine density in neurons expressing p.R206Q or p.R114W, whereas mature spine density was significantly elevated across all variants except p.R206Q **(Fig. 6S)**.

Together, these findings demonstrate that pathogenic hnRNP H2 variants disrupt structural hippocampal neuronal development by altering dendritic architecture and promoting premature or aberrant spine maturation *in vitro*, processes that may contribute to synaptic dysfunction in *HNRNPH2*-RNDD.

### Pathogenic hnRNP H2 variants impair cortical neuronal development *in vivo*

Having established that pathogenic hnRNP H2 variants disrupt dendritic architecture and spine maturation in cultured rat hippocampal neurons, we next asked whether these alterations also extend to neuronal development *in vivo*. To address this question, we used our previously validated molecular replacement system and delivered these constructs into mouse cortical progenitors by IUE at E15.5. Neuronal positioning and morphology were subsequently analysed at postnatal day 21 (P21) **(Fig. 7A)**. Analysis of cortical lamination revealed that neurons expressing the p.R114W variant exhibited a significant increase in the proportion of ectopically positioned GFP-positive cells within cortical layer VI compared with WT-expressing neurons, and reduced percentage of cells in cortical layer II/III. In contrast, the p.R206W, p.R206G, and p.R206Q variants displayed no detectable migration defects **(Fig. 7B-D)**. Although these latter variants reached the appropriate cortical layer, neurons expressing the p.R206G variant exhibited a significant increase in the proportion of misoriented neurons within the cortical plate, whereas p.R206W, p.R206Q, and p.R114W neurons were comparable to WT **(Fig. 7B, E, F)**. These findings suggest that distinct *HNRNPH2* variants differentially affect neuronal migration and post-migratory orientation.

**Figure 7.**
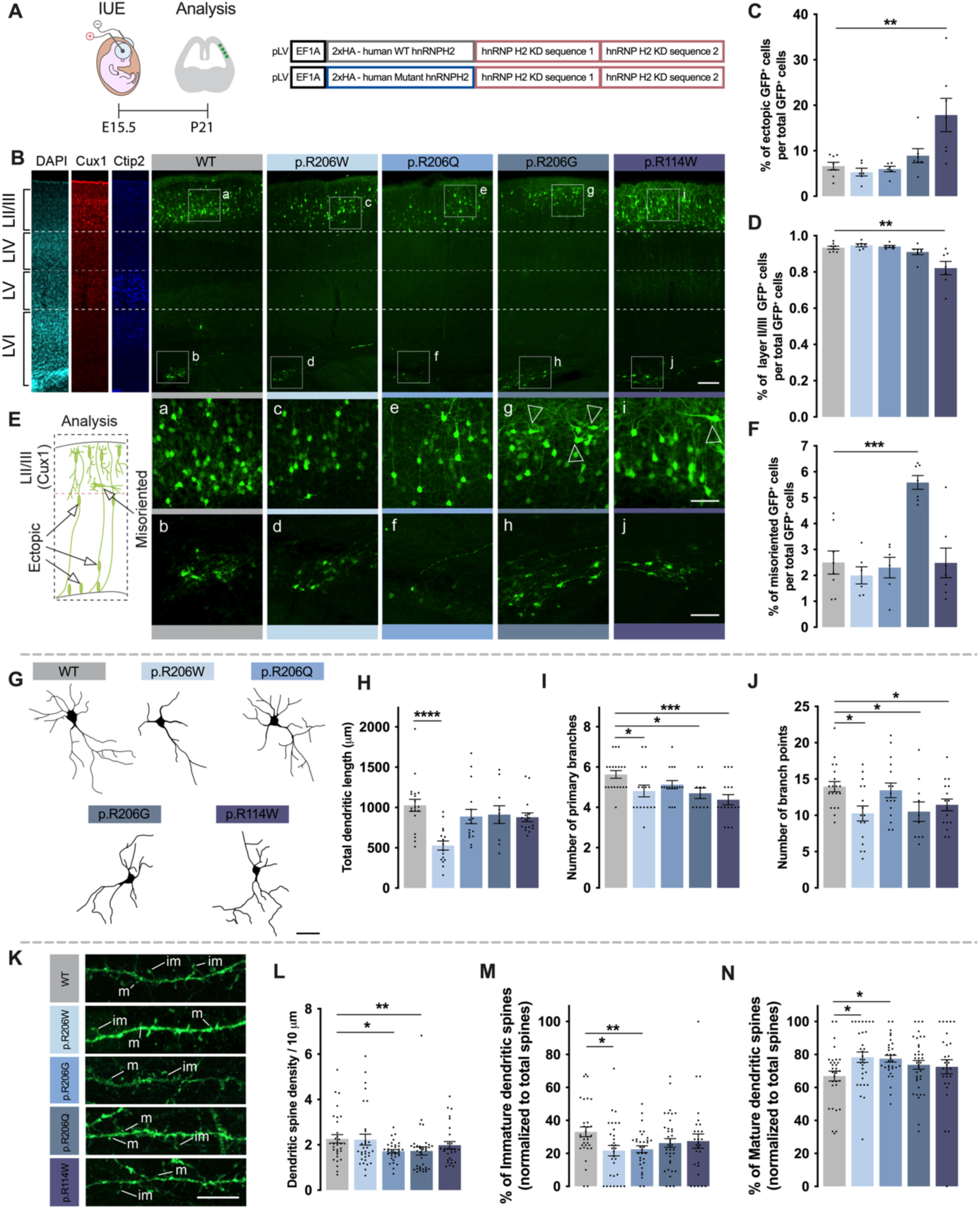
Disease-associated hnRNP H2 variants impair neuronal migration, dendritic development and dendritic spine maturation *in vivo*. **(A)** Experimental paradigm for the *in utero* analysis of neuronal migration, dendritic morphology and dendritic spine development. Mouse embryos were electroporated at embryonic day 15.5 (E15.5) with bicistronic lentiviral plasmids encoding HA-tagged human modified hnRNP H2 wild-type (WT) or disease-associated hnRNP H2 variants (p.R114W, p.R206Q, p.R206W or p.R206G), together with two knockdown sequences targeting endogenous rodent hnRNP H2. A separate GFP-expressing plasmid was co-electroporated to label electroporated neurons. Brains were collected at the indicated postnatal day 21 (P21), sectioned, and processed for confocal imaging. The schematic illustrates the IUE procedure targeting progenitors in the embryonic somatosensory cortex. **(B)** Representative confocal images of coronal cortical sections from brains expressing WT or mutant hnRNP H2. GFP-positive neurons (green) are shown together with DAPI (grey), Cux1 (red) and Ctip2 (blue). Insets highlight representative electroporated neurons. White arrowheads indicate misoriented neurons. **(C)** Quantification of the percentage of ectopic (mispositioned) GFP-positive cells (cortical layer VI) relative to the total number of GFP-positive cells in the same coronal section. **(D)** Quantification of the percentage of cortical layer II/III GFP-positive cells relative to the total number of GFP-positive cells in the same coronal section. **(E)** The schematic illustrates misoriented cells at cortical layer II/III and ectopic cells remaining in lower layers (cortical layer VI). **(F)** Quantification of the percentage of misoriented GFP-positive cells relative to the total number of GFP-positive cells in the same coronal section. **(G)** Representative reconstructions of electroporated cortical neurons used for morphological analysis. **(H–J)** Quantification of dendritic complexity in cortical neurons, including total dendritic length (H), number of primary branches (I), and number of branch points (J). **(K)** Representative images of dendritic segments from HA-tagged human hnRNP H2 WT or disease-associated hnRNP H2 variants (p.R114W, p.R206Q, p.R206W or p.R206G) neurons displaying multiple dendritic spines. m – mature, im – immature. **(L–N)** Quantification of dendritic spine morphology, including dendritic spine density per dendritic length (L), percentage of immature dendritic spines normalized to total number of spines (M) and percentage of mature dendritic spines normalized to total number of spines (N). Data are expressed relative to control and presented as mean ± SEM from 3 independent in utero electroporation experiments. At least 1 coronal brain section per animal was analysed for neuronal migration, and multiple reconstructed cortical neurons and dendritic segments were analysed per condition for dendritic morphology and spine analyses. Statistical significance was determined using unpaired two-tailed Student’s *t*-tests/Mann–Whitney tests where appropriate following assessment of data normality. p < 0.05 (*), p < 0.01 (**), p < 0.001 (***), p < 0.0001 (****); ns, not statistically significant. Scale bars: (B) 50 μm; inset, 15 μm; (G) 20 μm; (K) 10 μm.

We next examined whether pathogenic hnRNP H2 variants similarly affect dendritic maturation *in vivo* by reconstructing GFP-positive pyramidal neurons and quantifying dendritic complexity **(Fig. 7G)**. Neurons expressing the p.R206W variant exhibited a significant reduction in total dendritic length relative to WT, whereas p.R206G, p.R206Q, and p.R114W neurons were comparable to controls **(Fig. 7H)**. This was accompanied by a significant reduction in the number of primary dendrites in neurons expressing the p.R206W, p.R206Q and p.R114W variants, but not p.R206G **(Fig. 7I)**. Likewise, the total number of branch points was significantly reduced in neurons expressing the p.R206W, p.R206Q and p.R114W variants compared with WT, but not p.R206G **(Fig. 7J)**. Given the alterations observed in dendritic complexity, we next investigated the effect of disease-associated hnRNP H2 variants on cortical dendritic spine morphology *in vivo* **(Fig. 7K)**. Dendritic spine density was significantly reduced in cortical neurons expressing p.R206G and p.R206Q variants compared to WT-expressing neurons **(Fig. 7L)**. Classification of spines according to maturity revealed a significant reduction in the proportion of immature spines, accompanied by a corresponding increase in mature spines, in neurons expressing p.R206W and p.R206G compared to WT **(Fig. 7M, N)**.

Together, these findings extend our *in vitro* observations by demonstrating that pathogenic hnRNP H2 variants disrupt cortical neuronal development *in vivo*, with the overall pattern of dendritic and spine alterations broadly mirroring what we observed in cultured rat hippocampal neurons. Beyond variant-specific effects on neuronal migration and orientation, these variants impair dendritic maturation and alter dendritic spine maturation, indicating that hnRNP H2 dysfunction broadly compromises the developmental processes required for proper cortical circuit assembly and function.

### hnRNP H2 variants alter AMPAR-mediated excitatory synaptic transmission

Dysregulation of AMPAR-mediated synaptic transmission is a well-established hallmark of various neurodevelopmental disorders (Achuta et al., 2018; Ismail et al., 2022; Salpietro et al., 2019). Building on our findings that hnRNP H2 is required for maintaining excitatory synaptic strength, we next investigated whether disease-associated variants in hnRNP H2 disrupt AMPAR-mediated synaptic transmission.

We first assessed the surface and synaptic distribution of the GluA1 AMPA receptor subunit in hippocampal neurons expressing either WT or mutant hnRNP H2 **(Fig. 8A, B)**. Total surface GluA1 levels were significantly elevated in neurons expressing the p.R206W and p.R206G variants, with an upward trend observed in p.R114W-expressing neurons, although this did not reach statistical significance **(Fig. S7)**. At the postsynaptic level, this pattern was maintained: neurons expressing p.R206W and p.R206G hnRNP H2 exhibited a significant increase in the number, integrated density, and area of GluA1 puncta at synaptic sites, colocalising with PSD-95, while p.R114W-expressing neurons again showed a non-significant upward trend **(Fig. 8C-E)**. Despite alterations in overall PSD-95 levels across conditions **(Fig. 8F-H)**, the fraction of GluA1-positive PSD-95 puncta remained unchanged **(Fig. 8I)**. These findings suggest that while AMPAR abundance is increased at individual synapses in the mutant conditions, the overall number of GluA1-positive synapses is not altered.

**Figure 8.**
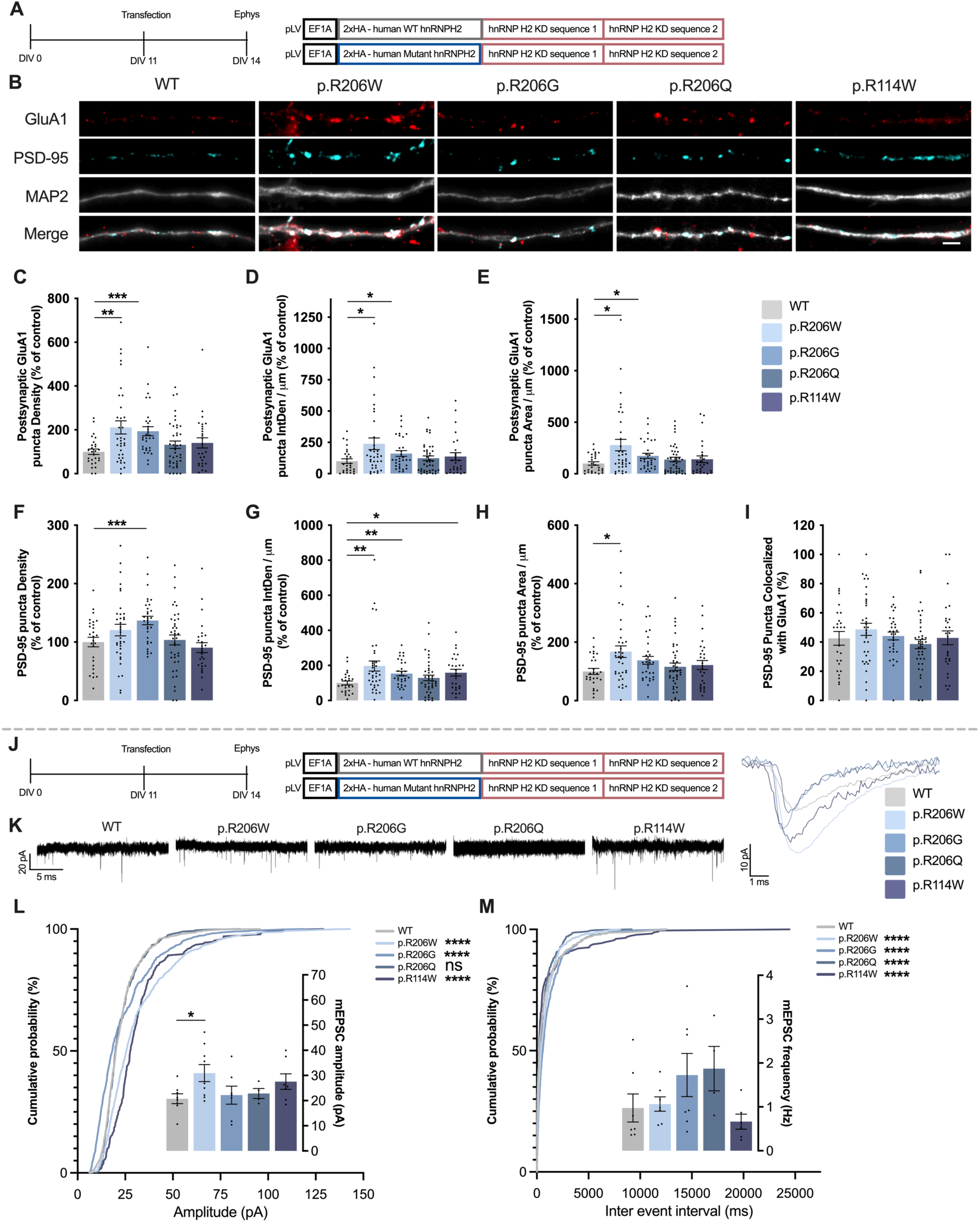
Disease-associated hnRNP H2 variants alter AMPA receptor-mediated synaptic transmission **(A)** Experimental paradigm for the analysis of AMPA receptor (AMPAR) surface expression. Primary rat hippocampal neurons were transfected at DIV 11 with bicistronic lentiviral plasmids expressing HA-tagged human modified hnRNP H2 wild-type (WT) or disease-associated hnRNP H2 variants (p.R114W, p.R206Q, p.R206W or p.R206G) together with two knockdown sequences targeting endogenous rodent hnRNP H2. Neurons were fixed at DIV 14 and immunostained for the AMPAR subunit GluA1, PSD-95 and MAP2. GFP was used to identify transfected neurons. **(B)** Representative images of dendritic segments from neurons expressing WT or mutant hnRNP H2 immunostained for surface GluA1 (red), PSD-95 (cyan) and MAP2 (grey). Merged images are shown. **(C–E)** Quantification of postsynaptic (PSD-95-positive) GluA1 surface puncta density (C), integrated density per dendritic length (D), and puncta area per dendritic length (F). **(F–I)** Quantification of PSD-95 puncta density (F), integrated density per dendritic length (G), and puncta area per dendritic length (H) and percentage of postsynaptic (PSD-95-positive) puncta containing surface GluA1 (I). **(J)** Experimental paradigm for electrophysiological recordings of AMPAR-mediated miniature excitatory postsynaptic currents (mEPSCs). Primary rat hippocampal neurons were transfected at DIV 11 with bicistronic lentiviral plasmids expressing HA-tagged human modified hnRNP H2 wild-type (WT) or disease-associated hnRNP H2 variants (p.R114W, p.R206Q, p.R206W or p.R206G) together with two knockdown sequences targeting endogenous rat hnRNP H2. A separate GFP-expressing plasmid was co-transfected to enhance neuronal morphology visualization. Whole-cell patch-clamp recordings were performed at DIV 14–15. AMPAR-mediated mEPSCs were pharmacologically isolated using tetrodotoxin (TTX), bicuculline (Bic) and D-AP5. **(K)** Representative traces of AMPAR-mediated mEPSCs (left) and averaged single mEPSC events (right) recorded from neurons expressing WT or mutant hnRNP H2. **(L, M)** Quantification of the cumulative probability distributions and mean values of mEPSC amplitude (L) and frequency (M). Data are expressed relative to control and presented as mean ± SEM from 3 independent neuronal preparations. AMPAR surface expression and synaptic immunocytochemistry analyses were performed in 3 independent neuronal preparations, with at least 12–15 fields of view containing multiple dendrites analysed per condition in each experiment. Electrophysiological recordings were obtained from at least 4 independent neuronal preparations. Statistical significance was determined using two-tailed Mann–Whitney tests for non-normally distributed datasets, two-tailed unpaired Student’s *t*-tests for normally distributed datasets, and Kolmogorov–Smirnov tests for cumulative probability distributions. p < 0.05 (*), p < 0.01 (**), p < 0.001 (***), p < 0.0001 (****); ns, not statistically significant. Scale bars: (B) 2.5 μm.

To determine whether these molecular changes in AMPAR distribution translate into functional alterations in synaptic transmission, we performed whole-cell patch-clamp recordings to measure AMPA receptor-mediated mEPSCs in transfected hippocampal neurons **(Fig. 8J, K)**.

Neurons expressing p.R206W showed a significant increase in mEPSC amplitude relative to WT, while p.R206G, p.R206Q and p.R114W were comparable to WT. The analysis of pooled mEPSC events confirmed a significant increase in amplitude for p.R206W, p.R206G and p.R114W **(Fig. 8L)**, consistent with the elevated GluA1 surface and synaptic levels observed by immunocytochemistry in these mutants **(Fig. 8L)**. In contrast, mEPSC frequency did not differ significantly across genotypes, despite the trends observed. However, the pooled event distribution revealed significant differences for p.R206W, p.R206G, p.R206Q, and p.R114W, when compared with WT **(Fig. 8M)**.

Together, these results demonstrate that pathogenic variants in hnRNP H2, particularly within the NLS domain, increase AMPAR-mediated synaptic transmission by raising postsynaptic receptor abundance, without substantially affecting presynaptic function

## Discussion

The molecular basis of *HNRNPH2*-related neurodevelopmental disorder has remained poorly understood, in part because the neuronal functions of hnRNP H2 had not been yet defined. RNA-binding proteins, including members of the hnRNP family, are critical for neuronal development and synaptic function (Lourinho et al., 2025), but the specific role of hnRNP H2 in these processes was unknown. Here we show that hnRNP H2 is enriched at synapses and is required for neuronal migration, dendritic arborization, spine maturation, and excitatory synaptic transmission. Pathogenic variants alter hnRNP H2 subcellular localisation, leading to defects in neuronal migration, dendritic and spine development both *in vitro* and *in vivo*, as well as impaired AMPA receptor-mediated currents. These findings establish hnRNP H2 as a key regulator of neuronal maturation and provide a mechanistic link between disease-associated variants and the neurodevelopmental phenotypes associated with *HNRNPH2*-RNDD.

hnRNP H2 has previously been shown to regulate the alternative splicing of TRF2, a telomere-binding protein whose truncated isoform (TRF2-S) promotes neuronal differentiation, by repressing TRF2-S production and thereby acting as a negative regulator of this process (Grammatikakis et al., 2016). More recent work indicates that co-depletion of hnRNP H1 and H2 impairs axonal mRNA translation, implicating these proteins in axonal regulation and local protein synthesis (S. J. Lee et al., 2018). However, the spatial, temporal, and mechanistic roles of hnRNP H2 during neuronal development had not been defined. In this work we show that hnRNP H2 is differentially expressed in developing hippocampal neurons, enriched in synaptic compartments, and recruited to synapses in an activity-dependent manner. Functional knockdown experiments reveal that hnRNP H2 is required for dendritic arborization, spine maturation, and excitatory synaptic transmission. *In vivo*, loss of hnRNP H2 perturbs cortical neuronal migration, suggesting an essential role during early circuit assembly and supporting a model in which hnRNP H2 links early events in cortical lamination with the later establishment of dendritic architecture and synaptic maturation. These findings indicate that hnRNP H2 is not confined to the nucleus but is a spatially dynamic RNA-binding protein whose coordinated distribution across nuclear, somatodendritic, and synaptic compartments enables it to influence multiple stages of neuronal development. A striking observation of our study is that even modest perturbations in the subcellular localisation of hnRNP H2 (such as those caused by mutations in its nuclear localisation signal - NLS) produce distinct and sometimes opposing effects on neuronal development. In cultured hippocampal neurons, at DIV14, hnRNP H2 knockdown enhances dendritic arborization but impairs spine maturation and synaptic transmission, whereas expression of NLS variants reduces dendritic arborization complexity while accelerating spine maturation and increasing synaptic transmission. These contrasting outcomes indicate that hnRNP H2’s developmental functions depend critically on the balance between its nuclear and cytoplasmic pools. As such, we propose that hnRNP H2 serves distinct roles across cellular compartments: in the nucleus, regulating RNA metabolism for transcripts such as TRF2 to influence global developmental programs; and in somatodendritic and synaptic compartments, controlling localised RNA metabolism, including regulation of GluA1 mRNA at synapses. This coupling of nuclear RNA regulation with local translational control may allow hnRNP H2 to coordinate large-scale developmental programs with fine-tuned synaptic modifications. Maintaining this balance likely depends on regulated nuclear import via Kapβ2, dendritic trafficking, and incorporation into RNA granules. Systematic mapping of hnRNP H2 distribution across neurodevelopment, combined with identification of its RNA targets, will be essential to define how this protein orchestrates RNA metabolism across neuronal compartments.

A central goal of this study was to define the functional consequences of pathogenic hnRNP H2 variants. Previous work reported that NLS variants accumulate in the cytoplasm due to impaired Kapβ2-mediated nuclear import (Gonzalez et al., 2023; Kreienkamp et al., 2022). Recently, it was demonstrated that NLS mutations in human iPSC-derived neurons and knock-in mice alter global gene expression and alternative splicing, further supporting the idea that these variants disrupt hnRNP H2 subcellular distribution and impair RNA metabolism (Korff et al., 2023). Collectively, these studies led to the proposal that NLS mutations cause a toxic gain of function through cytoplasmic accumulation, a mechanism consistent with our findings. Indeed, we show that NLS mutations lead to hnRNP H2 accumulation in the cytoplasm and within dendritic and synaptic compartments, likely through increased incorporation into RNA granules actively transported to synapses. This mislocalisation suggests pathogenic effects via dysregulated local RNA metabolism, drawing parallels to the amyotrophic lateral sclerosis (ALS)-associated RNA-binding protein FUS, in which cytoplasmic and synaptic accumulation contribute to disease through altered local RNA regulation (Sahadevan et al., 2021; Scekic-Zahirovic et al., 2016; Scekic-Zahirovic et al., 2021). Our data further suggest that mislocalised hnRNP H2 promotes enhanced local translation of specific RNA cargoes, with GluA1 emerging as a strong candidate. This hypothesis is further supported by evidence that GluA1 can be locally synthesized at synapses (Jung et al., 2020) and that GluA1 mRNA is present in dendrites of hippocampal neurons (Cox & Racca, 2013). Supporting this, neurons expressing NLS-mutant hnRNP H2 exhibit elevated synaptic GluA1-containing AMPA receptors, a change that could disrupt synaptic homeostasis. However, reduced nuclear hnRNP H2 in NLS-mutant neurons points to an additional loss-of-function mechanism. Given the predominantly nuclear localisation of hnRNP H2 under normal conditions, decreased nuclear localisation could impair critical RNA processing events, including splicing of TRF2 and potentially other neuronal transcripts. This model is analogous to TAR DNA-binding protein 43 (TDP-43)pathology in ALS and frontotemporal dementia (FTD), where nuclear depletion of this hnRNP family member leads to widespread splicing disruptions (Brown et al., 2022; E. Y. Liu et al., 2019; Ma et al., 2022; Melamed et al., 2019). We therefore propose a dual-pathogenic mechanism in which toxic cytoplasmic gain of function and loss of nuclear function act in parallel, potentially converging on shared downstream pathways.

Mutations outside the NLS can also disrupt hnRNP H2 function. The pathogenic p.R114W variant, with the mutation located in the second quasi-RNA recognition motif, has been shown to alter RNA metabolism and impair the interaction between hnRNP H2 and members of the large assembly of splicing regulators (LASR), a multimeric complex involved in alternative pre-mRNA splicing (Kreienkamp et al., 2022). We find that p.R114W increases hnRNP H2 conformational stability, reduces synaptic recruitment, and produces dendritic and synaptic phenotypes similar to those seen in NLS mutants. Although overall subcellular distribution is maintained, p.R114W likely disrupts hnRNP H2’s regulatory function in both nuclear and synaptic compartments by directly affecting its RNA-binding domain, thereby altering RNA target recognition and regulation. The similarity of phenotypes produced by NLS and qRRM variants, despite affecting distinct domains, suggests convergence on common RNA targets whose dysregulation drives disease. Mapping the physiological RNA targets of hnRNP H2 and defining how NLS and qRRM mutations alter their splicing, localisation, and translation in each compartment, will be essential for linking molecular defects to neuronal phenotypes.

Functionally, pathogenic hnRNP H2 variants consistently altered dendritic complexity and spine morphology, paralleling abnormalities reported across diverse models of neurodevelopmental disorders (G. L. Caldeira et al., 2022; Durand et al., 2012; Stuss et al., 2012). Surprisingly, a paradoxical and striking feature of our results is that most pathogenic variants accelerated aspects of synaptic development, yielding a higher density of morphologically mature dendritic spines and elevated synaptic AMPA receptor levels, hallmarks of premature synaptic maturation. This phenotype resembles early synaptic overdevelopment in some autism spectrum disorder (ASD) models, raising the possibility that hnRNP H2 variants trigger similar developmental programs (Forrest et al., 2018; X. Liu et al., 2016). Premature synapse maturation has been shown to compromise developmental windows for synaptic plasticity and circuit refinement, with long-term consequences for network function (Pires et al., 2026; Vermaercke et al., 2024). Although we did not directly assess synaptic plasticity, several observations point toward hnRNP H2 involvement in this process. First, we show that hnRNP H2 regulates basal synaptic AMPA receptor levels and is dynamically recruited to synapses in response to neuronal activity. Second, other hnRNP family members, most notably hnRNP A2/B1, are known to regulate activity-dependent local translation of GluA1 mRNA during synaptic plasticity (Jung et al., 2020). Thus, hnRNP H2 may perform a similar role, and its disruption could interfere with the fine-tuning of synaptic strength during synaptic plasticity. In the hippocampus, a region essential for learning and memory, such dysregulation could impair cognitive function; indeed, male mice harbouring the p.R206W variant in *HNRNPH2* exhibit deficits in spatial learning and memory (Korff et al., 2023). Future studies assessing activity-dependent plasticity in neurons expressing hnRNP H2 variants will be critical to link these molecular disruptions to cognitive phenotypes in *HNRNPH2*-RNDD.

Although primarily based on *in vitro* findings, our results also highlight a plausible therapeutic avenue for *HNRNPH2*-RNDD. A common effect of most disease-associated hnRNP H2 variants is elevated synaptic GluA1-containing AMPA receptor levels. Dysregulated AMPAR signaling is increasingly recognized as a convergent mechanism across diverse neurodevelopmental disorders: mutations in genes encoding AMPAR subunits are linked to intellectual disability and developmental abnormalities, while aberrant AMPAR expression is observed in Fragile X syndrome, Rett syndrome, and in the cerebellum of individuals with ASD (Achuta et al., 2018; Ismail et al., 2022; Li et al., 2016; Purcell et al., 2001; Salpietro et al., 2019). This suggests that AMPAR dysregulation may underlie core pathophysiology in *HNRNPH2*-RNDD. Supporting this notion, studies in the valproic acid mouse model of autism demonstrate that increased AMPAR expression and heightened excitatory transmission coincide with behavioral abnormalities, which can be rescued by pharmacological AMPAR antagonism (Kim et al., 2019). By analogy, reducing AMPAR overactivity in hnRNP H2-mutant neurons may mitigate premature synapse maturation and excessive excitatory drive. Clinically approved negative AMPAR modulators, such as the anti-epileptic Perampanel, offer a readily translatable starting point (Ceolin et al., 2012). Whether modulating AMPAR function can reverse *HNRNPH2*-RNDD phenotypes remains to be tested, but knock-in mouse models carrying patient-derived NLS mutations now provide an experimentally tractable platform to address this question (Korff et al., 2023). More broadly, therapeutic strategies for *HNRNPH2*-RNDD are beginning to emerge. Recent preclinical work demonstrated that non-allele-specific antisense oligonucleotide therapy targeting murine *Hnrnph2* and thus reducing expression from both wild-type and mutant alleles, rescued molecular, neuropathological, and behavioral phenotypes in these knock-in mouse models (Korff et al., 2026). Together with the mechanistic insights presented here, these findings underscore a growing and increasingly tractable path toward treatment for *HNRNPH2*-RNDD and may inform therapeutic strategies for other neurodevelopmental disorders in which AMPAR dysregulation is a shared pathophysiological feature.

## Material and methods

### Animals

Wistar rats were used for the preparation of primary hippocampal neuronal cultures, and C57BL/6J mice (Charles River) were used for in utero electroporation experiments. Male and female animals were used throughout the study. Animals were maintained under standard housing conditions, and all procedures were conducted in accordance with Portuguese legislation (Decree-Law 113/2013) and the European Union Directive 2010/63/EU on the protection of animals used for scientific purposes. All animal procedures were approved by the Animal Experimentation and Welfare Committee of the University of Valencia and CNC-UC/CIBB.

### Primary cultures of rat hippocampal neurons

Primary hippocampal neuronal cultures were prepared from embryonic day 17–18 (E17–18) Wistar rat embryos, as previously described (De Luca et al., 2026). Pregnant females were anesthetized with isoflurane (IsoVet) and sacrificed by decapitation. Embryonic hippocampi were dissected and transferred to ice-cold Ca²⁺- and Mg²⁺-free Hank’s balanced salt solution (HBSS) containing 5.36 mM KCl, 0.44 mM KH₂PO₄, 137 mM NaCl, 4.16 mM NaHCO₃, 0.34 mM Na₂HPO₄·2H₂O, 5 mM glucose, 1 mM pyruvic acid, 10 mM HEPES and 0.001% phenol red (pH 7.2). Under sterile conditions, tissue was incubated in 0.06% trypsin in HBSS for 15 min at 37 °C, and enzymatic digestion was terminated with HBSS containing 10% fetal bovine serum. Hippocampi were washed twice with HBSS and transferred to Neurobasal medium supplemented with SM1 (1:50; STEMCELL Technologies), 25 µM glutamate, 0.5 mM glutamine and 0.12 mg/mL gentamycin (Gibco, Thermo Fisher Scientific). Tissue was mechanically dissociated and the cell suspension was passed through a 70-µm cell strainer. Cell density was determined using trypan blue exclusion and a hemocytometer.

For low-density cultures, neurons were plated at 8.8 × 10³ cells/cm² onto 18-mm glass coverslips previously coated with poly-D-lysine (PDL; 0.1 mg/mL in 0.166 M boric acid, pH 8.2). Cells were allowed to adhere for 2 h at 37 °C in a humidified atmosphere containing 5% CO₂/95% air, after which coverslips were transferred to dishes containing cultured cerebrocortical glial cells. Neuronal and glial layers were physically separated by paraffin dots placed on the neuronal side of the coverslips. For western blot, neurons were plated at 90 × 10³ cells/cm² onto multi-wells.

At day in vitro 3 (DIV 3), 10 µM 5-fluoro-2ʹ-deoxyuridine (FDU) was added to inhibit glial proliferation. Neuronal cultures were maintained in a humidified incubator with 5% CO2 / 95% air, at 37 °C, for 7 to 21 days, depending on the experimental approaches. Every 7 days, one third of the culture medium was replaced with fresh medium of identical composition but lacking glutamate.

### Molecular modelling and molecular dynamics (MD) simulations

A structural model of human hnRNP H2 was generated by homology modelling using the hnRNP H2 amino acid sequence, UniProt accession P55795 (The UniProt Consortium, 2017), as the target. The structure of the highly homologous hnRNP H1 protein, PDB ID: 6DHS), https://www.rcsb.org/structure/6DHS, (Penumutchu et al., 2018), which shares 96% sequence identity with hnRNP H2, was selected as the template. The template structure was prepared using PyMOL v3.0.4 by retaining chain A and removing the remaining chains and non-relevant molecules. Target and template sequences were aligned using ClustalW (Thompson et al., 1994) with the BLOSUM substitution matrix. Residues 1–220 of hnRNP H2 were modelled, encompassing the qRRM1 and qRRM2 RNA-binding domains and the region containing the nuclear localisation signal (NLS), where several *HNRNPH2*-NDD-associated variants are located. Homology modelling was performed using MODELLER v10.5, (Eswar et al., 2006) generating 100 candidate structures. The final model was selected based on its Discrete Optimized Protein Energy (DOPE) score (Shen & Sali, 2006) and structural quality assessment using ProSA-web (Wiederstein & Sippl, 2007). The selected model was subsequently aligned with the template in PyMOL, and structural similarity was evaluated by calculating the root-mean-square deviation (RMSD) between the Cα atoms of all modelled residues and the corresponding Cα atoms of the template. The wild-type (WT) hnRNP H2 model and models carrying the *HNRNPH2*-NDD-associated variants p.R114W, p.R206Q, p.R206W, and p.R206G were subsequently prepared for molecular dynamics (MD) simulations using CHARMM-GUI (Jo et al., 2008; J. Lee et al., 2016). These variants were selected to investigate structural perturbations affecting functionally distinct regions of hnRNP H2: p.R114W is located within the qRRM2 RNA-binding domain, whereas substitutions at p.R206 occur within the NLS-containing region. Each system was prepared at pH 7.0, the protonation states of ionizable residues were determined using PyPka software (Reis et al., 2024), and solvated in a rectangular box containing TIP3P water molecules. Na⁺ and Cl⁻ ions were added to neutralize the system and establish an ionic concentration of 0.15 M. Periodic boundary conditions were applied, and long-range electrostatic interactions were calculated using the Particle-Mesh Ewald (PME) method. The CHARMM36 force field (Huang & MacKerell, 2013) was used, and the topology, coordinate, minimization, equilibration, and production input files generated by CHARMM-GUI were used to perform the simulations in GROMACS 2019.4 (Abraham et al., 2015).

Prior to MD simulations, each system was subjected to energy minimization using the steepest-descent algorithm for a maximum of 50,000 steps to remove steric clashes and unfavorable atomic contacts. The minimized systems were subsequently equilibrated before production MD simulations. The first stage was conducted under constant-volume and constant-temperature conditions using the NVT ensemble, whereas the subsequent seven stages were conducted under constant-pressure and constant-temperature conditions using the NPT ensemble at 310 K and 1 bar. Temperature was controlled using a velocity-rescale thermostat, and pressure was controlled using the Parrinello– Rahman barostat during the NPT stages. Positional restraints on the protein backbone and side chains, together with dihedral restraints, were progressively reduced throughout the equilibration procedure. Production simulations were performed under constant-pressure and constant-temperature conditions using the NPT ensemble at 303.15 K. Three independent 500-ns replicas were generated for WT hnRNP H2 and each of the four variants, corresponding to an aggregate simulation time of 1.5 µs per system and 7.5 µs across all five systems.

To assess mutation-associated conformational changes, the relative positioning of the first two qRRM domains was monitored throughout the trajectories. The interdomain distance was defined between residues 11 and 130, located within the structured regions of qRRM1 and qRRM2, respectively. In parallel, the relative orientation of these domains was evaluated using the angle defined by residues 11, 90, and 130, with residue 90 located in the linker connecting qRRM1 and qRRM2. Distance–angle distributions were monitored as a function of simulation time, and joint distance–angle distributions were generated across the three 500-ns replicas to characterize the conformational space sampled by WT and mutant proteins. The dispersion and temporal evolution of these measurements were used to compare differences in interdomain orientation, compactness, conformational flexibility, and stability among WT and variant proteins.

In addition, representative mutant structures were aligned with a representative structure from the WT simulations using the conserved qRRM1 domain as the reference region. Structural comparisons were performed using PyMOL to examine mutation-associated changes in the orientation of qRRM2 and the NLS-containing region and to complement the quantitative distance–angle analysis. Together, these analyses were designed to determine whether *HNRNPH2*-NDD-associated variants alter the conformational dynamics of regions implicated in RNA recognition and nucleocytoplasmic localisation.

### hnRNP H2 knockdown and molecular replacement strategy

A miR30-based shRNA strategy was used to silence endogenous rodent *Hnrnph2* mRNA. For knockdown experiments, the hnRNP H2 knockdown construct co-expressed GFP, enabling identification and morphological analysis of transfected neurons. A matched GFP-expressing vector lacking the hnRNP H2-targeting sequence was used as control. The *Hnrnph2*-targeting silencing sequence used in these experiments (hereafter referred to as hnRNP H2 KD sequence 1) was 5ʹ-CATTATAACCTCCGTAGTCATC-3ʹ.

For molecular replacement experiments, bicistronic lentiviral plasmids were designed to simultaneously deplete endogenous rodent *Hnrnph2* mRNA and express an shRNA-resistant human hnRNP H2 transgene (NM_001032393.3), thereby functionally replacing endogenous hnRNP H2, allowing the functional consequences of WT and disease-associated variants to be assessed. The constructs contained two miR30-based silencing sequences targeting rodent *Hnrnph2*, selected based on the rat *Hnrnph2* reference sequence and subsequently adapted to a miR30-based shRNA architecture: hnRNP H2 silencing sequence 1 (5ʹ-CATTATAACCTCCGTAGTCATC-3ʹ) and hnRNP H2 silencing sequence 2 (5ʹ-TTTAAAGCCAATTTCACTTCAT-3ʹ). Thus, endogenous rodent *Hnrnph2* mRNA was silenced by the two miR30-based shRNAs, while human hnRNP H2 was expressed in the same cells to functionally replace the depleted endogenous protein. Given the high sequence identity between human and rodent hnRNP H2 (∼99%), synonymous substitutions were introduced into the human coding sequence at the silencing sequence target sites to prevent recognition and silencing of the replacement transgene while preserving the hnRNP H2 amino acid sequence (nt234 AAA>AAG, nt240 GCT>GCA, nt241 TTG>CTG, nt777 TAT>TAC, and nt780 AAT>AAC). This shRNA-resistant human sequence therefore served as the WT replacement construct and as the backbone for generating the disease-associated variants. A Kozak consensus sequence and two tandem HA epitopes were incorporated at the N-terminus, and the resulting modified WT hnRNP H2 sequence was cloned into the BamHI/XbaI sites of the pLV-EF1α backbone, upstream of the miR30-based shRNA cassettes.

Disease-associated hnRNP H2 variants used in this study included the NLS variants p.R206W, p.R206G, and p.R206Q, and the qRRM variant p.R114W. Mutant constructs were generated from the codon-optimized human WT hnRNP H2 construct by site-directed mutagenesis using mutation-specific primers and the Q5 Site-Directed Mutagenesis Kit (New England Biolabs, #E0554S), according to the manufacturer’s instructions. The following nucleotide substitutions were introduced: CGG><u>T</u>GG for p.R206W, CGG><u>G</u>GG for p.R206G, CGG>C<u>A</u>G for p.R206Q, and CGG><u>T</u>GG for p.R114W.

Plasmids were amplified in competent Escherichia coli DH5α cells. Briefly, cells were transformed with 1 µg plasmid DNA by incubation on ice for 10 min, heat shock at 42 °C for 45 s and recovery on ice for 5 min. Bacteria were plated on LB-agar containing ampicillin and incubated for 16 h at 37 °C. Individual colonies were expanded in LB medium containing ampicillin, and plasmid DNA was purified using the HiPure Plasmid Filter Maxiprep Kit (Invitrogen). DNA concentration and purity were assessed using a NanoDrop 2000c spectrophotometer (Thermo Scientific), and constructs were verified by DNA sequencing before experimental use.

### Western blotting

Total protein extracts were prepared from hippocampal cultures at DIV 7-21. Cells were washed twice with ice-cold PBS, and then lysed in RIPA buffer containing 150 mM NaCl, 50 mM Tris-HCl, 5 mM EGTA, 1% Triton X-100, 0.5% deoxycholic acid, and 0.1% sodium dodecyl sulfate (SDS), supplemented with a protease inhibitor cocktail comprising 0.1 mM PMSF, 1 μg/mL chymostatin, 1 μg/mL leupeptin, 1 μg/mL antipain, and 1 μg/mL pepstatin (Sigma-Aldrich), together with 50 mM NaF and 1.5 mM sodium orthovanadate. Lysates were frozen at −80 °C, thawed, and centrifuged at 16,000 × g for 10 min. Protein concentration in the resulting supernatants was determined using the bicinchoninic acid (BCA) assay (Thermo Scientific), according to the manufacturer’s instructions.

Samples were diluted in Laemmli sample buffer (Bio-Rad) supplemented with 5% β-mercaptoethanol and denatured for 5 min at 95 °C. Proteins were separated by SDS-PAGE using 4-20% Mini-PROTEAN TCX polyacrylamide gels (Bio-Rad, #4561094) for approximately 2 h at 80 V and subsequently transferred onto polyvinylidene fluoride (PVDF) membranes (Millipore) for 75 min at 110 V. Membranes were blocked in EveryBlot Blocking buffer (Bio-Rad, #12010020) and incubated overnight at 4 °C with primary antibodies diluted in blocking buffer. Membrane was washed 3 times for 10 min with TBS-T containing 20 mM Tris, 150mM NaCl, and 0.1% Tween, pH 7.6.

Following washing, membranes were incubated for 1 h at room temperature with StarBright Blue fluorophore-conjugated secondary antibodies: goat anti-rabbit IgG StarBright Blue 700 (Bio-Rad, #12004162) and goat anti-mouse IgG StarBright Blue 520 (Bio-Rad, #12004159), both diluted 1:5,000 in blocking buffer. hnRNP H2 was detected using an anti-hnRNP H2 antibody (rabbit, 1:1000; Abcam, ab179439), while anti-β-actin antibody (mouse 1:200,000; Sigma-Aldrich, A5441) was used as a loading control. Fluorescence signals were acquired using a ChemiDoc MP Imaging System (Bio-Rad), and band immunoreactivity was quantified using ImageJ software (National Institutes of Health).

### Induction of synaptic activity

To induce excitatory synaptic activity, at DIV 14/15 hippocampal neurons were exposed for 30 min to conditioned culture medium containing 50 µM bicuculline, 2.5 mM 4-aminopyridine (4-AP) and 10 µM glycine, adapted with minor modifications from (M. V. Caldeira et al., 2013). Control cultures received an equivalent volume of conditioned medium. As the drugs were dissolved in water, control cells received an equivalent volume of water. Immediately following stimulation, cells were fixed and processed for immunocytochemistry.

### Calcium-phosphate transfection

Primary hippocampal neurons cultured on 18-mm coverslips were transfected at DIV 4 or DIV 10-11, depending on the experimental endpoint, using a calcium phosphate precipitation protocol adapted with minor modifications from (Afonso et al., 2019). Plasmid DNA was diluted in TE buffer containing 10 mM Tris-HCl and 2.5 mM EDTA (pH 7.3). For transfections with a single construct, 1.5 µg of the plasmid of interest was used per coverslip. When co-transfection was required, 1.5 µg of the plasmid of interest was combined with 0.5 µg of a GFP-expressing plasmid, yielding a total of 2 µg DNA per coverslip. A 2.5 M CaCl₂ solution prepared in 10 mM HEPES was then added dropwise to the DNA solution, and the resulting mixture was added dropwise to 2× HEPES-buffered saline (HEBS; 274 mM NaCl, 10 mM KCl, 1.4 mM Na₂HPO₄, 11 mM dextrose, and 42 mM HEPES, pH 7.2).

DNA precipitates were added to neuronal cultures maintained in conditioned Neurobasal medium previously supplemented with 2 mM kynurenic acid, and cells were incubated for 2 h at 37 °C. Cultures were subsequently washed for 20 min with Neurobasal medium containing 2 mM kynurenic acid and lightly acidified with HCl, followed by a further 20-min incubation. The medium was then replaced with the original conditioned culture medium, and neurons were maintained in a humidified incubator with 5% CO2 / 95% air, at 37 °C, for 72 h to allow transgene expression.

Neurons analysed at DIV7 were transfected at DIV4, whereas experiments performed at DIV_14/15 used neurons transfected at DIV10-11. This timing was used for both knockdown and molecular replacement experiments.

### Immunocytochemistry

At the indicated DIV in the figures, neurons were fixed for 15 min in 4% paraformaldehyde containing 4% sucrose in PBS, washed three times with PBS and permeabilized with 0.25% Triton X-100 in PBS for 5 min. Following washing, cells were blocked with 3% BSA in PBS for 30 min at room temperature (RT). Primary antibodies were diluted in blocking solution, centrifuged at 16,600 × g for 20 min at 4 °C and incubated with neurons overnight at 4 °C. Cells were subsequently washed three times with PBS and incubated with the appropriate secondary antibodies for 1 h, at RT. After secondary antibody incubation, cells were washed twice with 0.1% Triton X-100 in PBS followed by PBS and mounted using Dako fluorescent mounting medium.

Primary antibodies included anti-GFP (mouse, 1:100; Roche, 11814460001), anti-GluA1 (rabbit, 1:80; Alomone Labs, AGC-004), anti-HA (rabbit, 1:500; Cell Signaling Technology, #3724), anti-hnRNP H2 (rabbit, 1:250; Abcam, ab179439), anti-Homer1 (guinea pig, 1:500; Synaptic Systems, 160 004), anti-MAP2 (chicken, 1:100; Abcam, ab5392), anti-PSD-95 (mouse, 1:200; Invitrogen, MA1-045; Guinea pig, 1:200; SySy, 124014)) and anti-vGluT1 (guinea pig, 1:100; Sigma-Aldrich, AB5905). Alexa Fluor-conjugated secondary antibodies were used at 1:500.

### Surface GluA1 live staining

Surface GluA1-containing AMPA receptors were labelled in live DIV14 hippocampal neurons by incubation with an antibody directed against the extracellular domain of GluA1, diluted in conditioned culture medium, for 10 min at RT, adapted with minor modifications from (Afonso et al., 2019; De Luca et al., 2026). Neurons were rapidly washed three times with PBS and fixed with 4% PFA for 15 min. Following fixation, cells were washed three times with PBS for 5 min each and incubated with the appropriate secondary antibody, diluted in 3% BSA in PBS, for 1 h at RT to selectively label surface GluA1 before membrane permeabilization. Cells were subsequently washed three times with PBS for 5 min each and permeabilized with 0.25% Triton X-100 in PBS for 5 min. Following permeabilization, neurons were processed for immunocytochemistry as described above, including blocking and immunolabelling for the additional intracellular markers.

### *In utero* electroporation

In utero electroporation (IUE) of the embryonic somatosensory cortex was performed at embryonic day 15.5 (E15.5) following previously described procedures (Szczurkowska et al., 2016), with minor modifications. Briefly, E15.5 timed-pregnant C57BL/6J mice were anaesthetized with isoflurane (2.5% for induction and 1.5% during surgery), and the uterine horns were exposed by laparotomy. Plasmid DNA (1.5 μg/μL) and pCAGGs-IRES-GFP (0.5 μg/μL) were diluted in water and combined with Fast Green (0.3 mg/mL; Sigma) to facilitate visualization of the injection.

For hnRNP H2 knockdown experiments, embryos were electroporated with either or the previously described GFP-expressing control vector (Ctr) or hnRNP H2 knockdown construct, which co-expressed GFP to enable identification of electroporated neurons. For experiments assessing disease-associated hnRNP H2 variants, embryos were electroporated with the previously described bicistronic molecular replacement constructs encoding HA-tagged human hnRNP H2 WT or the disease-associated variants p.R114W, p.R206Q, p.R206W, or p.R206G, together with two knockdown sequences targeting endogenous rodent *Hnrnph2*. In these experiments, a separate pCAGGs-IRES-GFP plasmid was co-electroporated with the hnRNP H2 constructs to label electroporated neurons.

Approximately 5–6 μL of the DNA solution was injected unilaterally through the uterine wall into the lateral ventricle of each embryo. Following injection, the embryo’s head was positioned between 5-mm tweezer-type circular electrodes to target progenitor cells in the developing somatosensory cortex. Six electrical pulses (30 V, 50-ms duration, 950-ms intervals) were delivered using a square-wave electroporation generator (ECM 830; BTX Harvard Apparatus). Only one cerebral hemisphere per embryo was electroporated. Left and right hemispheres were alternated across experimental conditions to minimize potential lateralization bias.

Following electroporation, the uterine horns were returned to the abdominal cavity, and embryos were allowed to continue normal development. For hnRNP H2 knockdown experiments assessing neuronal migration, brains were collected at E18. For experiments assessing the effects of disease-associated hnRNP H2 variants on neuronal positioning, dendritic morphology, and dendritic spine development, brains were collected at P21 and processed for subsequent histological and imaging analyses.

### Tissue collection

All brains were fixed by transcardial perfusion with 4% paraformaldehyde in PBS, cryopreserved in 30% sucrose overnight and sectioned coronally (80 μm thickness) for immunostaining or (20 μm thickness) for immunohistochemistry using a vibrotome (Leica VT1200S) or cryostat (Leica CM1900) respectively.

### Immunohistochemistry

Immunohistochemistry was performed as previously described (Calvo Parra et al., 2025), with minor adaptations. Briefly, free-floating brain sections were permeabilized and blocked in PBS containing 0.2% Triton X-100 and 10% horse serum (HS). Sections were incubated overnight at 4 °C with primary antibodies against Cux1 (rabbit, 1:500; Proteintech, 11733-1-AP), Ctip2 (rat, 1:250; Abcam, ab18465), and GFP (chicken, 1:500; Aves Labs), diluted in PBS containing 0.2% Triton X-100 and 1% HS. Cux1 and Ctip2 immunostaining was used to delineate cortical layers, while GFP immunostaining was used to identify neurons labelled by in utero electroporation.

Following primary antibody incubation, sections were washed in PBS and incubated for 2 h at room temperature with the appropriate fluorescently conjugated secondary antibodies: anti-chicken Alexa Fluor 488 (1:700; Jackson ImmunoResearch), anti-rat Alexa Fluor 647 (1:400; Jackson ImmunoResearch), and anti-rabbit Alexa Fluor 555 (1:400; Jackson ImmunoResearch), diluted in PBS containing 0.2% Triton X-100 and 1% HS. Sections were counterstained with DAPI (1:5000; Sigma-Aldrich) and mounted using FluorSave Reagent (Millipore) for subsequent fluorescence and confocal microscopy.

### Widefield, confocal and super-resolution microscopy

For quantitative fluorescence analysis of primary neuronal cultures, images were acquired using a Plan-Apochromat 63× oil-immersion objective (NA 1.40) on an inverted Axio Observer.Z1/7 microscope equipped with an AxioCam HR R3 camera and ZEN Blue software (Carl Zeiss). Within each experiment, acquisition parameters, including exposure time and fluorescence illumination intensity, were kept constant across experimental conditions. Random fields of view encompassing somatic or dendritic regions were acquired, and image analyses were performed blind to experimental condition.

For high-resolution characterization of hnRNP H2 synaptic localisation, primary hippocampal neurons were fixed and immunostained for hnRNP H2 together with MAP2 and the presynaptic marker vGluT1 and/or the postsynaptic markers Homer1 and PSD-95. DIV21 neurons were imaged using an LSM 980 confocal microscope equipped with Airyscan 2 (Carl Zeiss). Images were processed using Airyscan processing and Airyscan joint deconvolution, and projections were generated using ZEN Blue software. Three-dimensional reconstruction and visualization of dendritic segments were performed using Imaris software.

For super-resolution analysis, DIV 14 neurons were imaged using an Elyra 7 microscope equipped with Lattice SIM² super-resolution technology (Carl Zeiss). Super-resolution images were used to examine the spatial distribution of hnRNP H2 relative to vGluT1-positive presynaptic and Homer1- or PSD-95-positive postsynaptic compartments.

For *in vivo* analyses, immunofluorescence images of brain sections containing GFP⁺ neurons labelled by in utero electroporation were acquired using an Olympus Fluoview FV10i confocal microscope. Images for neuronal migration analysis were acquired using a 10× objective, whereas images for dendritic morphology analysis were acquired using a 10× objective with a digital zoom of 40×. For dendritic spine analysis, high-resolution images of GFP⁺ neurons were acquired using a Zeiss LSM 980 confocal microscope equipped with a 40× water-immersion objective.

### Quantitative fluorescence and colocalisation analysis

For quantitative fluorescence analysis, cells from the experimental conditions being compared were cultured, processed, and immunostained in parallel and imaged using identical acquisition settings. Images from randomly selected fields of view encompassing somatic or dendritic regions were analysed, depending on the experimental endpoint. Fluorescent puncta corresponding to proteins of interest and their colocalisation with synaptic markers were quantified using an in-house macro developed for ImageJ/Fiji (National Institutes of Health). For dendritic analyses, regions of interest (ROIs) were randomly selected based on MAP2 immunostaining, which was also used to determine dendritic length.

For each protein of interest, an appropriate fluorescence intensity threshold was established to identify recognizable puncta above background, and comparable threshold levels were maintained across experimental conditions within each experiment. Particle analysis was then performed to determine puncta number (density), integrated density (IntDen) and total puncta area. Measurements were normalized to dendritic length, unless otherwise specified in the corresponding figure legends.

For colocalisation analyses, overlap with the respective postsynaptic marker was used to identify synaptically localised puncta. Homer1-positive puncta and PSD-95-positive puncta were used to assess postsynaptic localisation of hnRNP H2 and surface GluA1. Synaptic localisation was quantified as the percentage of postsynaptic marker-positive puncta containing the protein of interest. In addition, the density, integrated density, and area of colocalised puncta were quantified and normalized to dendritic length, where applicable. All image analyses were performed blind to experimental condition.

### Analysis of cortical neuronal migration

For analysis of cortical migration, immunofluorescence images were acquired using an Olympus Fluoview FV10i confocal microscope with a 10× objective. GFP-positive neurons were quantified according to their position and orientation within defined cortical regions/layers.

### Dendritic morphology and Sholl analysis

In vitro, dendritic arborization was analysed from images acquired with a Plan-Apochromat 20×/0.8 objective on an Axio Observer.Z1/7 microscope (Carl Zeiss). Within each experiment, acquisition parameters, including exposure time and fluorescence intensity, were kept constant across experimental conditions. MAP2 immunostaining was used as a somatodendritic marker to delineate neuronal morphology, while GFP or HA immunoreactivity, depending on the experimental paradigm, was used to identify transfected neurons. Neuronal morphology was quantified using the Simple Neurite Tracer (SNT) plugin in ImageJ/Fiji (National Institutes of Health). Sholl analysis was performed using concentric circles centred on the soma and spaced at 5-µm radial intervals. The number of dendritic intersections as a function of distance from the soma, total dendritic length, number of primary dendrites, branch points, and terminal branches were determined for each neuron. Soma area was additionally quantified where indicated. Analyses were performed blind to experimental condition, preferentially using isolated neurons.

For *in vivo* postnatal morphological analysis, GFP⁺ cortical neurons labelled by in utero electroporation (IUE) at E15.5 were analysed at P21. Confocal z-stacks were maximum-intensity projected in Fiji, and individual GFP⁺ neurons were reconstructed for quantitative analysis following previously described procedures (Calvo Parra et al., 2025). Dendritic morphology was assessed from neuronal reconstructions, and total dendritic length, number of primary branches, and number of branch points were quantified for each neuron. Analyses were performed at the single-neuron level and compared across experimental conditions.

### Dendritic spine analysis

For dendritic spine analysis, secondary dendritic segments were imaged using a Plan-Apochromat 63× oil-immersion objective (NA 1.40, Oil DIC M27) on an LSM 710 Axio Examiner confocal microscope (Carl Zeiss) (equipped with ZEN Black 2012 software. Within each experiment, acquisition parameters, including exposure time and fluorescence intensity, were kept constant across experimental conditions. Transfected neurons were identified by eGFP fluorescence (AF480 channel). Images were acquired from secondary dendrites within a 29.99 × 29.99 µm field of view. Z-stacks were acquired using a constant z-step across experimental conditions, while the number of optical sections was adjusted as needed to encompass the full depth of each dendritic segment and ensure visualization of all dendritic spines. Image stacks were subsequently analysed using Imaris x64 9.6.1 (Bitplane AG), allowing three-dimensional reconstruction and visualization of dendritic segments for manual spine classification.

Dendritic spines were classified into five morphological categories, according to predefined morphological criteria (filopodia, branched, thin, stubby, and mushroom). Filopodia were defined as protrusions lacking a discernible head, whereas branched spines were defined as protrusions containing more than one head. Thin spines exhibited a clearly defined neck and a head diameter <0.5 µm, with a head width less than twice the neck width. Stubby spines lacked a clearly defined neck, whereas mushroom spines displayed a distinct neck and a head diameter ≥0.5 µm, with a head width greater than twice the neck width. For analyses of spine maturation, filopodia were classified as immature, whereas thin, stubby, and mushroom spines were classified as mature. Branched spines were considered a transitional morphology and were therefore excluded from the immature/mature classification, in accordance with previously established criteria (Pchitskaya et al., 2023). All image analyses and morphological classifications were performed blind to experimental condition.

For postnatal *in vivo* spine analysis, GFP⁺ cortical neurons labelled by in utero electroporation at E15.5 were analysed at P21. High-resolution images of GFP⁺ dendritic segments were acquired using a Zeiss LSM 980 confocal microscope equipped with a 40× water-immersion objective. GFP fluorescence was used to identify electroporated neurons and visualize dendritic segments. Dendritic spine density was quantified as the number of spines per dendritic length. Spine maturation was assessed by classifying dendritic spines as immature or mature based on their morphology, and the percentage of each category was calculated relative to the total number of spines analysed. Multiple dendritic segments from GFP⁺ cortical neurons were analysed per experimental condition, and analyses were performed blind to experimental condition.

### Whole-cell patch-clamp recordings

Whole-cell patch-clamp recordings were performed in cultured hippocampal neurons visualized under phase contrast using a Zeiss Axio Observer A1 inverted microscope equipped with a 40× oil-immersion objective (NA 1.4). Neurons were continuously perfused at approximately 2 mL/min with extracellular solution containing 140 mM NaCl, 10 mM glucose, 2.4 mM KCl, 4 mM MgCl₂, 4 mM CaCl₂ and 10 mM HEPES (pH 7.3; 300–310 mOsm). Recording pipettes were pulled from borosilicate glass capillaries (Science Products, GmbH) and had resistances of 3.5–5.5 MΩ when filled with intracellular solution containing 145 mM potassium gluconate, 10 mM HEPES, 1 mM EGTA, 2 mM MgCl₂·6H₂O, 2 mM MgATP and 0.3 mM NaGTP (pH 7.3; 300 mOsm).

AMPAR-mediated miniature excitatory postsynaptic currents (mEPSCs) were recorded at a holding potential of −70 mV for 5 min. The extracellular solution was supplemented with 500 nM tetrodotoxin (TTX), 10 µM bicuculline and 50 µM D-AP5 to block voltage-gated sodium channels, GABAA receptors and NMDA receptors, respectively. Recordings were acquired with a HEKA EPC10 USB amplifier using PatchMaster (HEKA Eletronik), filtered at 2.9 kHz and sampled at 10 kHz. Traces were analysed in Clampfit (Molecular Devices) using a template-matching routine. The first 50 detected events from each recording were used to determine mEPSC amplitude and frequency. The template mEPSC was obtained by manual selection, automatic alignment, and averaging of visually identified synaptic events. Analyses were performed blind to experimental condition.

### Statistical analysis

Data are presented as mean ± SEM from at least 3 independent experiments performed using independent biological preparations, as indicated in the corresponding figure legend. Where applicable, values from each independent experiment were normalized to the mean of the respective control condition and expressed relative to control. For microscopy experiments, at least 12 fields of view or neurons per experimental condition were generally analysed in each independent experiment, unless otherwise specified. All quantifications were performed blind to experimental condition.

Statistical analyses were performed using GraphPad Prism 10. Where applicable, data distribution was assessed for normality before statistical testing. Comparisons between two groups were performed using two-tailed unpaired Student’s *t*-tests for normally distributed data or two-tailed Mann–Whitney tests for non-normally distributed data, as appropriate. For Sholl analyses, statistical significance was assessed using two-way ANOVA followed by Sidak’s multiple comparisons test. For activity-dependent synaptic recruitment experiments involving multiple experimental conditions, two-way ANOVA followed by Tukey’s multiple comparisons test was used. Cumulative probability distributions were compared using the Kolmogorov–Smirnov test. The statistical test used for each experiment is specified in the corresponding figure legend. Statistical significance was defined as p < 0.05.

## Acknowledgements

Fluorescence imaging was performed at the MICC Imaging facility of CNC-UC, partially supported by PPBI – Portuguese Platform of BioImaging (PPBI-POCI-01-0145-FEDER-022122).

## Author contributions

A.L. and M.S.S. contributed to methodology, investigation, formal analysis, data curation and visualization. E.T., M.D. and A.D.P. contributed to methodology, investigation and formal analysis. S.C.P. contributed to methodology, investigation, formal analysis, data curation and visualization. G.D.P. and G.G. contributed to methodology, investigation and formal analysis. A.B.C. contributed to methodology, investigation, formal analysis, data curation, visualization and writing – original draft. L.P.A. contributed to conceptualization and methodology. P.P. contributed to methodology and writing – review and editing. I.S.M. contributed to conceptualization, methodology, formal analysis, writing – review and editing and supervision. R.J.N. contributed to conceptualization, methodology and writing – review and editing. M.M. contributed to methodology, investigation, writing – review and editing, supervision and project administration. A.W.C. contributed to conceptualization, methodology, investigation, formal analysis, data curation, visualization, writing – review and editing, supervision, project administration and funding acquisition. C.B.D. contributed to conceptualization, writing – review and editing and supervision. R.O.C. contributed to conceptualization, methodology, investigation, formal analysis, data curation, visualization, writing – review and editing, supervision, project administration and funding acquisition. All authors reviewed and approved the final version.

## Funding

This work was supported by Portuguese national funds through the Foundation for Science and Technology (Fundação para a Ciência e a Tecnologia – FCT), under the projects COMPETE2030-FEDER-00676600_RARE-TARGETS (project 15671), UIDP/04539/2025, and LA/P/0058/2020. Individual funding was provided to Rui O. Costa through DL57/2016/CP1448/CT0009, to Miranda Mele through 2020.00420-CEECIND, and to Emanuel Tahiri through UI/BD/154292/2022. This work was also supported by the Portuguese League Against Epilepsy (Liga Portuguesa Contra a Epilepsia – LPCE) through the Scientific Incentive Grant 2022 awarded to Rui O. Costa. In addition, this work was supported by the Generalitat Valenciana through Project No. CDEIGENT/2021/005, awarded to Andrzej W. Cwetsch.

## Data availability

The data supporting the findings of this study are available from the corresponding author upon reasonable request.

## Ethics declaration

Wistar rats were used for primary hippocampal neuronal cultures and C57BL/6J mice (Charles River) for in utero electroporation experiments. Male and female animals were used throughout the study. Animals were maintained under standard housing conditions, and all procedures were conducted in accordance with Portuguese legislation (Decree-Law 113/2013) and EU Directive 2010/63/EU. All animal procedures were approved by the Animal Welfare Body of the Centre for Neuroscience and Cell Biology, University of Coimbra/CIBB (ORBEA-CNC) and the Animal Experimentation and Welfare Committee of the University of Valencia. The principles of the 3Rs (Replacement, Reduction and Refinement) were applied throughout the study.

## Competing interests

The authors report no competing interests.

---

**Supplementary Figure S1.**
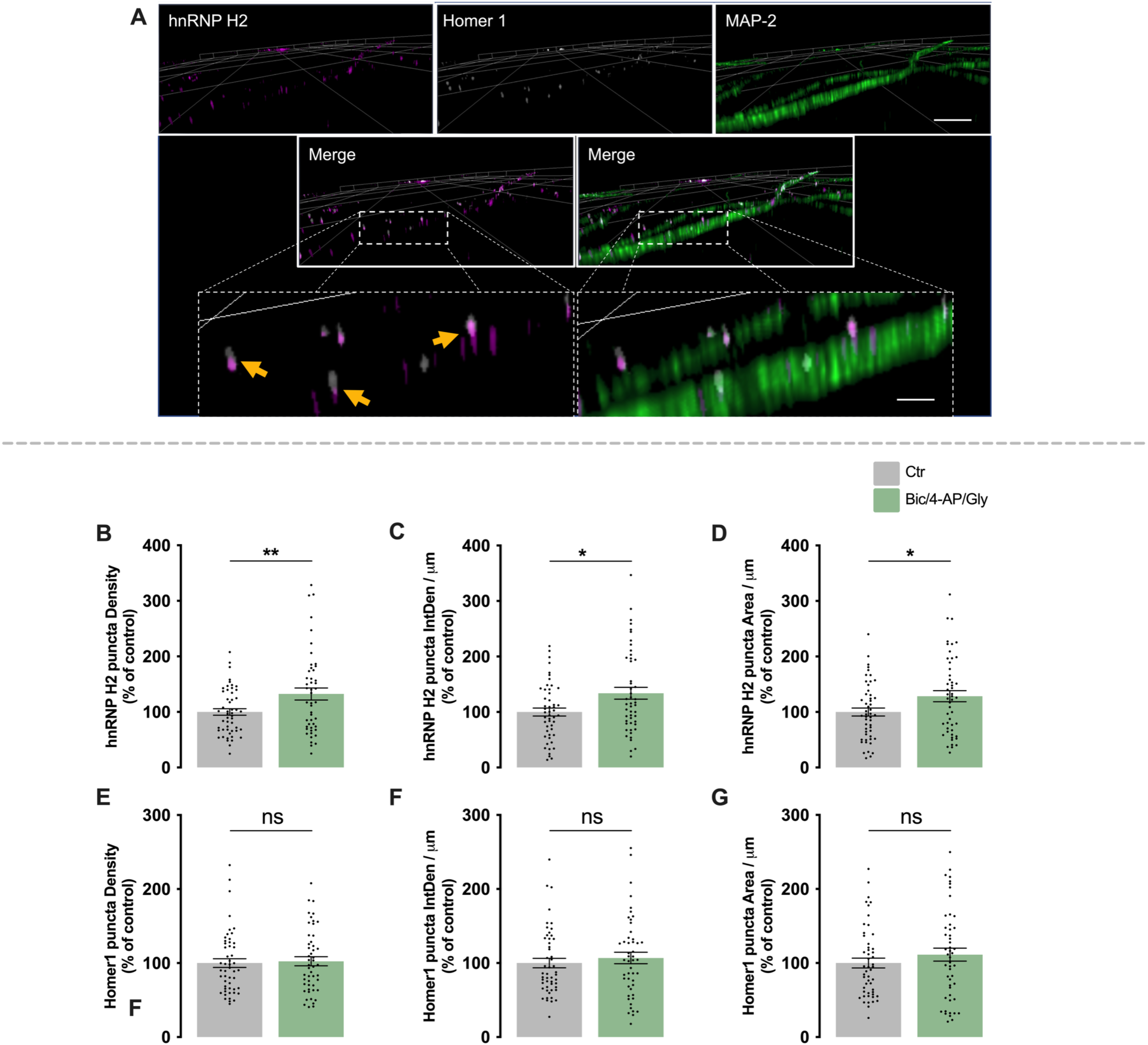
hnRNP H2 localises to excitatory synapses and is recruited in an activity-dependent manner in developing hippocampal neurons. Additional quantification from the experiments shown in Figure 1. **(A)** Representative projection of confocal images of DIV 21 hippocampal neurons immunostained for hnRNP H2 (magenta), Homer1 (grey) and MAP2 (green). Insets show hnRNP H2 puncta localised adjacent to Homer1-positive postsynaptic sites, at the dendritic spine neck, immediately below Homer1 along dendrites (arrows). The projection was performed with Zeiss ZEN blue software. **(B–D)** Quantification of the percentage of dendritic hnRNP H2 puncta density (B), integrated density per dendritic length (C), and puncta area per dendritic length (D). **(E–G)** Quantification of the percentage of dendritic Homer1 puncta density (B), integrated density per dendritic length (C), and puncta area per dendritic length (D). Data are expressed relative to control and presented as mean ± SEM from 4 independent neuronal preparations for immunocytochemistry analysis. At least 12 fields containing multiple dendrites were analysed per condition in each experiment. Statistical significance was determined using two-tailed unpaired Student’s *t*-tests. p < 0.05 (*), p < 0.01 (**); ns, not statistically significant. Scale bars: (B) 20 μm; insert 5 μm.

**Supplementary Figure S2.**
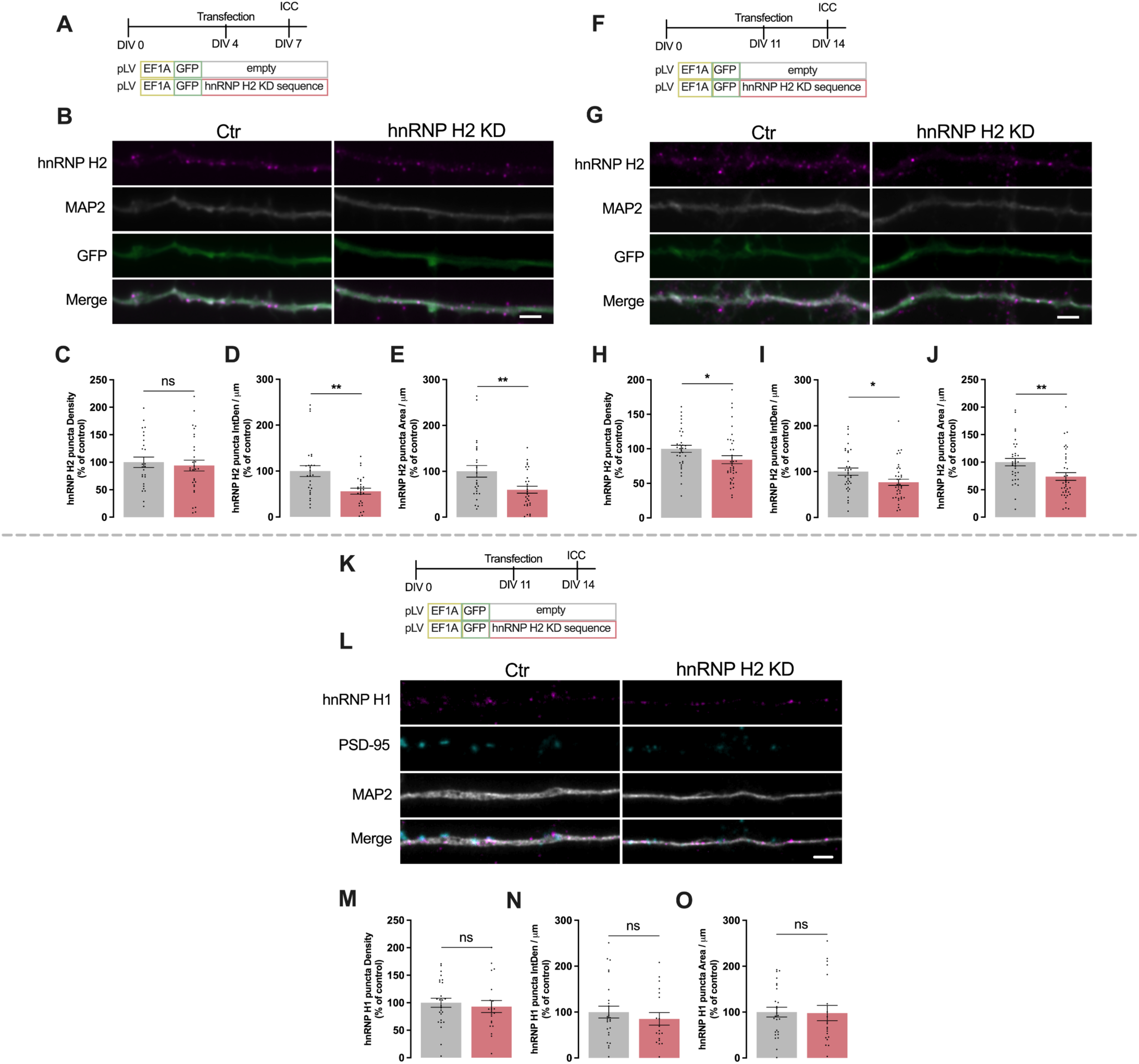
Validation of hnRNP H2 knockdown efficiency in primary hippocampal neurons. **(A)** Experimental paradigm for the analysis of the effect of an hnRNP H2-targeting knockdown sequence (hnRNP H2 KD sequence) in total hnRNP H2 in developing hippocampal neurons. Primary rat hippocampal neurons were transfected at DIV 4 with either a control plasmid (Ctr) or a hnRNP H2 KD sequence. Neurons were fixed at DIV 7 and immunostained for hnRNP H2 and MAP2, while GFP identified transfected neurons. **(B)** Representative images of dendritic segments from Ctr and hnRNP H2-knockdown neurons immunostained for hnRNP H2 (magenta), MAP2 (grey). GFP was used to identify transfected neurons. **(C–E)** Quantification of hnRNP H2 puncta density (C), integrated density per dendritic length (D), and puncta area per dendritic length (E). **(F)** Experimental paradigm for the analysis of the effect of an hnRNP H2-targeting knockdown sequence (hnRNP H2 KD sequence) in total hnRNP H2 in developing hippocampal neurons. Primary rat hippocampal neurons were transfected at DIV 11 with either a control plasmid (Ctr) or a hnRNP H2 KD sequence. Neurons were fixed at DIV 14 and immunostained for hnRNP H2 and MAP2, while GFP identified transfected neurons. **(G)** Representative images of dendritic segments from Ctr and hnRNP H2-knockdown neurons immunostained for hnRNP H2 (magenta), MAP2 (grey). GFP was used to identify transfected neurons. **(H–J)** Quantification of hnRNP H2 puncta density (H), integrated density per dendritic length (I), and puncta area per dendritic length (J). **(K)** Experimental paradigm for the analysis of the effect of an hnRNP H2-targeting knockdown sequence (hnRNP H2 KD sequence) in total hnRNP H1 in developing hippocampal neurons. Primary rat hippocampal neurons were transfected at DIV 11 with either a control plasmid (Ctr) or a hnRNP H2 KD sequence. Neurons were fixed at DIV 14 and immunostained for hnRNP H2 and MAP2, while GFP identified transfected neurons. **(L)** Representative images of dendritic segments from Ctr and hnRNP H2-knockdown neurons immunostained for hnRNP H1 (magenta), PSD/95 MAP2 (Cyan) and MAP2 (grey). **(M–O)** Quantification of hnRNP H1 puncta density (M), integrated density per dendritic length (N), and puncta area per dendritic length (O). Data are expressed relative to control and presented as mean ± SEM from 3 independent neuronal preparations. At least 12 neurons or 12 fields of view containing multiple dendrites were analysed per condition in each independent experiment. Statistical significance was determined using two-tailed unpaired Student’s *t*-tests. p < 0.05 (*), p < 0.01; ns, not statistically significant. Scale bars: (B, G, L) 2.5 μm;

**Supplementary Figure S3.**
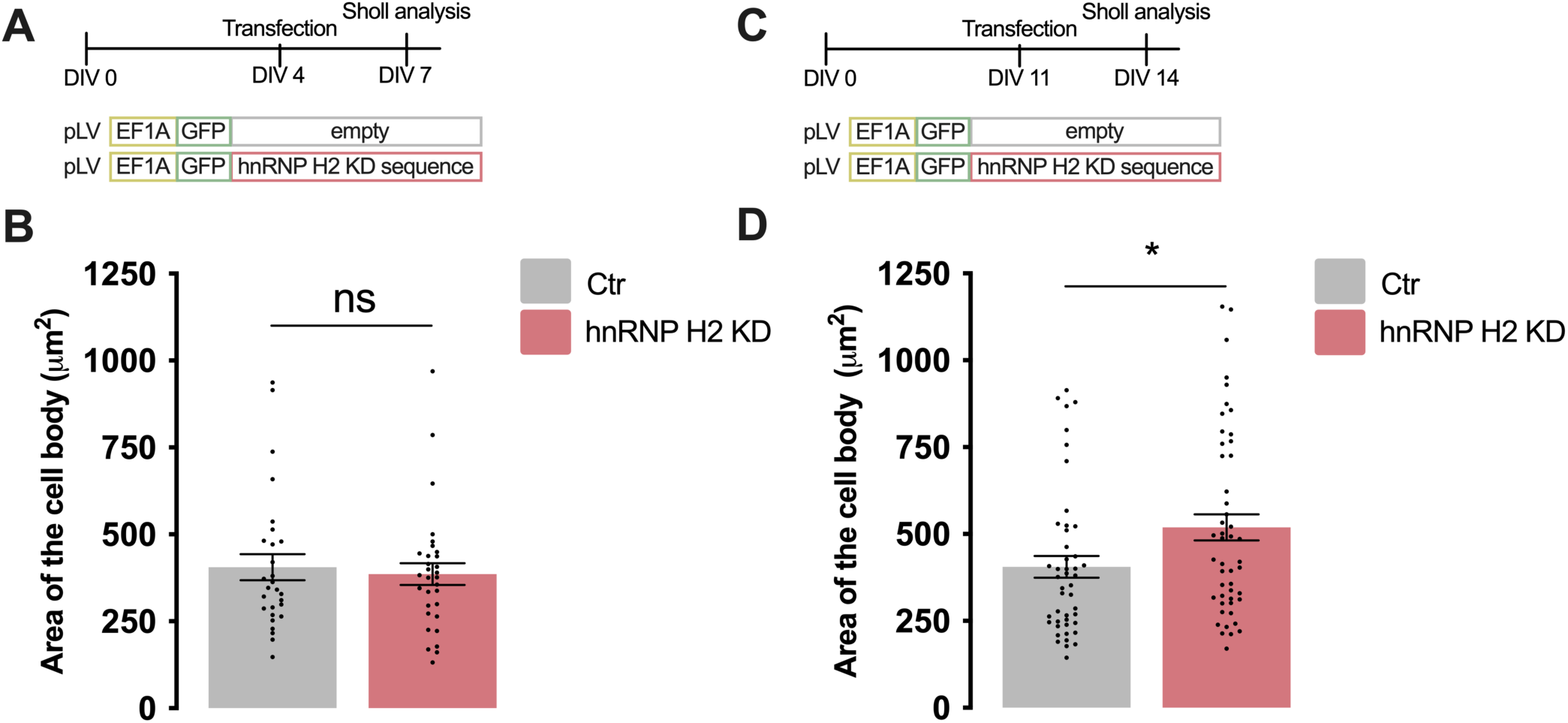
hnRNP H2 knockdown induces structural alterations in developing and mature hippocampal neurons. Additional quantification from the experiments shown in Figure 2. **(A)** Experimental paradigm for the analysis of neuronal soma area in developing hippocampal neurons. Primary rat hippocampal neurons were transfected at DIV 4 with either a control plasmid (Ctr) or a plasmid expressing an hnRNP H2-targeting knockdown sequence (hnRNP H2 KD sequence). Neurons were fixed at DIV 7 and immunostained for MAP2, while GFP identified transfected neurons. **(B)** Quantification of neuronal soma area. **(C)** Experimental paradigm for the analysis of neuronal soma area in mature hippocampal neurons. Primary rat hippocampal neurons were transfected at DIV 11 with either the Ctr or hnRNP H2 KD sequence plasmid and fixed at DIV 14 for MAP2 immunostaining. **(D)** Quantification of neuronal soma area. Data are presented as mean ± SEM from 3–4 independent neuronal preparations. At least 12 neurons per condition, in each experiment, were analysed. Data distribution was first assessed for normality. Normally distributed datasets were analysed using two-tailed unpaired Student’s *t*-tests, whereas non-normally distributed datasets were analysed using two-tailed Mann–Whitney tests. p < 0.05 (*); ns, not statistically significant.

**Supplementary Figure S4.**
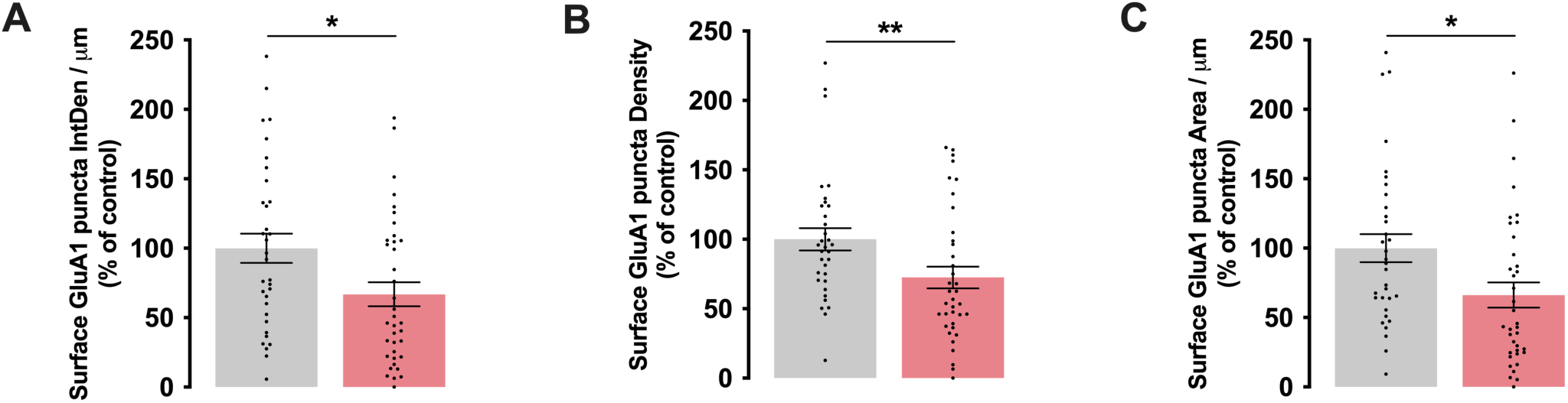
hnRNP H2 knockdown reduces AMPA receptor surface expression. Additional quantification from the experiments shown in Figure 4. **(A–C)** Quantification of the percentage of dendritic surface GluA1 puncta density (A), integrated density per dendritic length (B), and puncta area per dendritic length (C). Data are expressed relative to control and presented as mean ± SEM from 3 independent neuronal preparations. Statistical significance was determined using two-tailed Mann–Whitney tests. p < 0.05 (*), p < 0.01 (**).

**Supplementary Figure S5.**
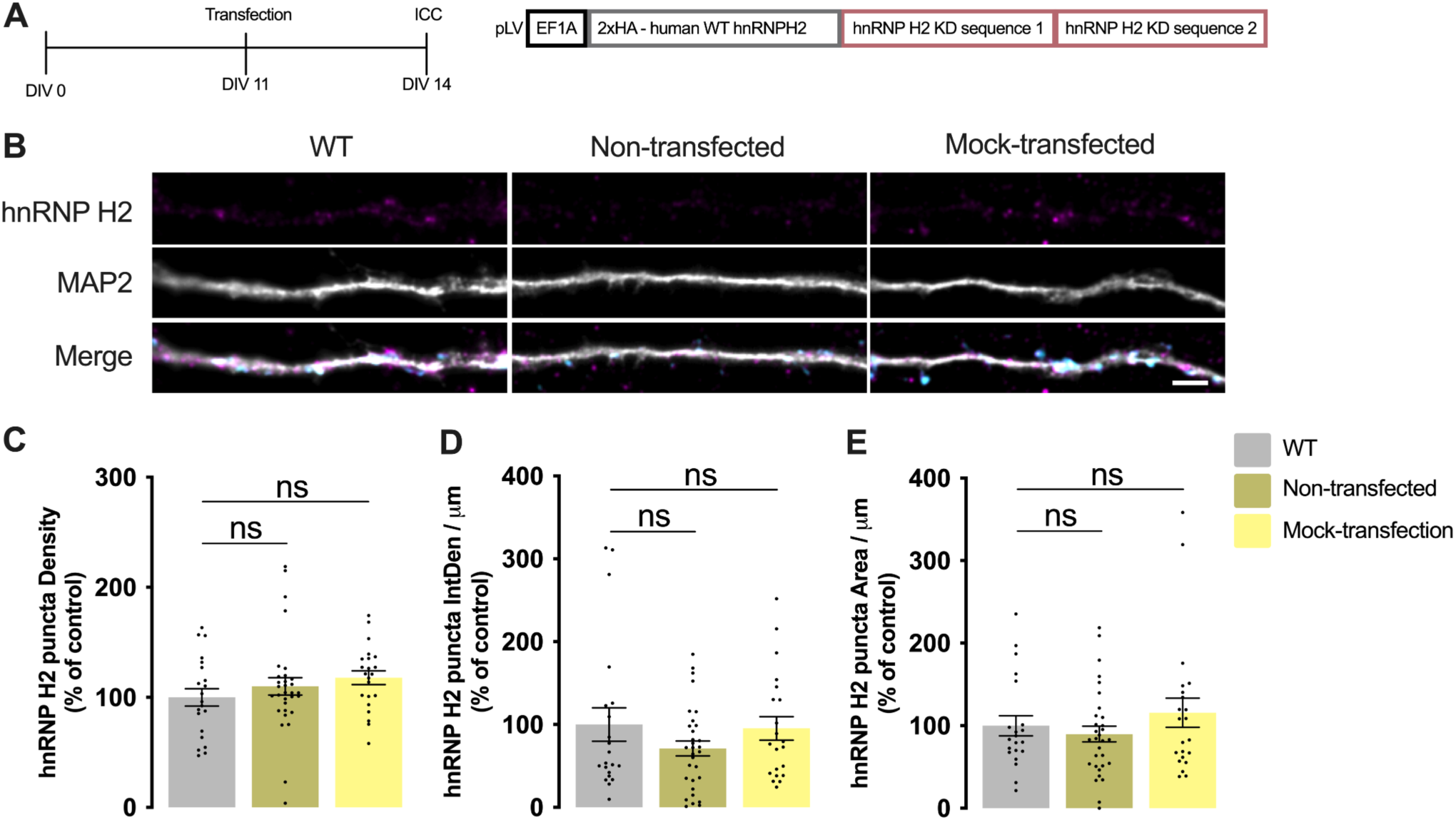
Molecular replacement preserves physiological levels of hnRNP H2 in rat hippocampal neurons. **(A)** Experimental paradigm for the analysis of total hnRNP H2 levels in hippocampal neurons following the molecular replacement strategy. Primary rat hippocampal neurons were either left non-transfected or transfected at DIV 11 with a bicistronic lentiviral plasmid encoding HA-tagged human hnRNP H2 wild-type (WT) together with two knockdown sequences targeting endogenous rodent hnRNP H2. Mock-transfected neurons were subjected to the transfection procedure without plasmid DNA. Neurons were fixed at DIV 14 and immunostained for hnRNP H2 and MAP2. **(B)** Representative images of dendritic segments from primary rat hippocampal neurons transfected with a bicistronic lentiviral plasmid encoding HA-tagged human hnRNP H2 wild-type (WT) together with two knockdown sequences targeting endogenous rodent hnRNP H2, non-transfected neurons, and mock-transfected neurons immunostained for hnRNP H2 (magenta) and MAP2 (grey). Merged images are shown. **(C–E)** Quantification of hnRNP H2 puncta density (C), integrated fluorescence intensity per dendritic length (D), and puncta area per dendritic length (E) in WT-transfected, non-transfected, and mock-transfected neurons. Data are expressed relative to control and presented as mean ± SEM from 3 independent neuronal preparations. Statistical significance was determined using two-tailed Mann–Whitney tests. ns, not statistically significant. Scale bar: (A) 2.5 μm.

**Supplementary Figure S6.**
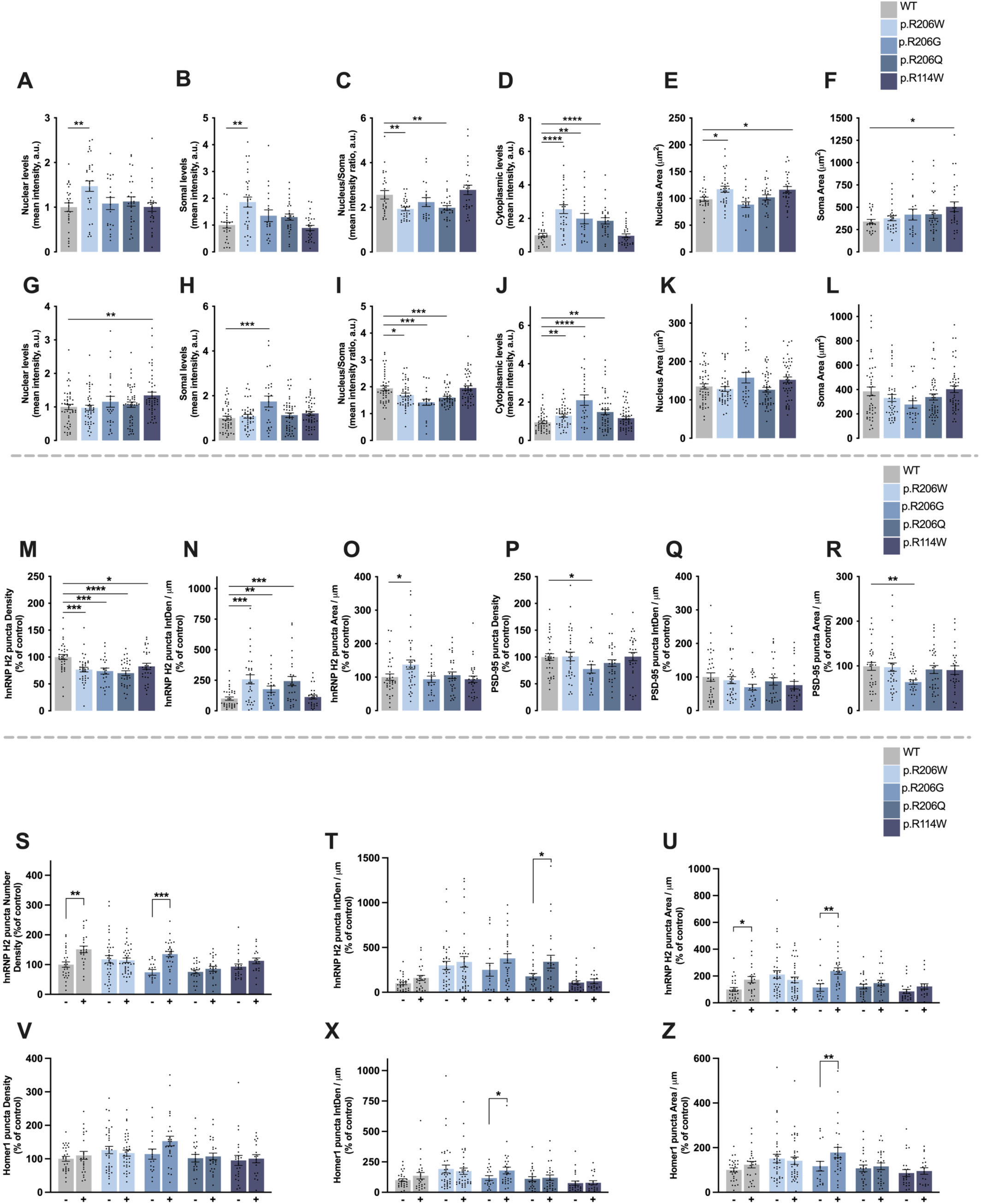
Disease-associated HNRNPH2 variants disrupt hnRNP H2 subcellular localisation and activity-dependent synaptic recruitment. Additional quantification from the experiments shown in Figure 5. **(A–F)** Quantification of hnRNP H2 nuclear intensity levels (A), somal intensity levels (B), nucleus/soma intensity ratio (C), cytoplasmic intensity levels (D), nuclear area (E) and soma area (F) in developing hippocampal neurons (DIV 7) expressing bicistronic lentiviral plasmids encoding HA-tagged human hnRNP H2 wild-type (WT) or the disease-associated variants p.R114W, p.R206Q, p.R206W or p.R206G, together with two knockdown sequences targeting endogenous rodent hnRNP H2. **(G–L)** Quantification of hnRNP H2 nuclear intensity levels (G), somal intensity levels (H), nucleus/soma intensity ratio (I), cytoplasmic intensity levels (J), nuclear area (K) and soma area (L) in mature hippocampal neurons (DIV 14) expressing bicistronic lentiviral plasmids encoding HA-tagged human hnRNP H2 wild-type (WT) or the disease-associated variants p.R114W, p.R206Q, p.R206W or p.R206G, together with two knockdown sequences targeting endogenous rodent hnRNP H2. **(M–O)** Quantification of dendritic hnRNP H2 puncta density (M), integrated density per dendritic length (N) and puncta area per dendritic length (O). **(P–R)** Quantification of PSD-95 puncta density (P), integrated density per dendritic length (Q) and puncta area per dendritic length (R). **(S–U)** Quantification of dendritic hnRNP H2 puncta density (S), integrated density per dendritic length (T) and puncta area per dendritic length (U) under basal and stimulated conditions with bicuculline (50 μM), 4-aminopyridine (2.5 mM) and glycine (10 μM) for 30 min before fixation. **(V–Z)** Quantification of Homer1 puncta density (V), integrated density per dendritic length (X) and puncta area per dendritic length (Z) under basal and stimulated conditions with bicuculline (50 μM), 4-aminopyridine (2.5 mM) and glycine (10 μM) for 30 min before fixation. Data are expressed relative to control and presented as mean ± SEM from 3 independent neuronal preparations for localisation analyses and 4 independent neuronal preparations for activity-dependent recruitment experiments. At least 12 neurons or 12 fields of view containing multiple dendrites were analysed per condition in each independent experiment. Statistical significance was determined using two-tailed unpaired Student’s *t*-tests for localisation analyses and two-way ANOVA followed by Tukey’s multiple comparisons test for activity-dependent recruitment. p < 0.05 (*), p < 0.01 (**), p < 0.001 (***), p < 0.0001 (****).

**Supplementary Figure S7.**
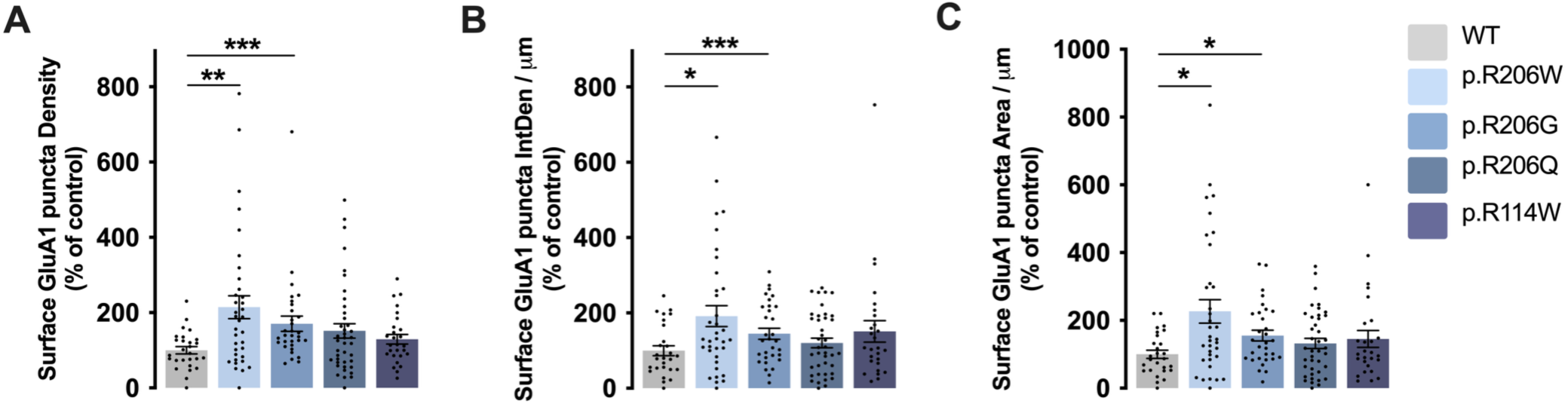
Disease-associated hnRNP H2 variants differentially affect surface AMPA receptor expression. Additional quantification from the experiments shown in Figure 8. **(A–C)** Quantification of surface GluA1 puncta density (A), integrated density per dendritic length (B), and puncta area per dendritic length (C) in primary rat hippocampal neurons expressing bicistronic lentiviral plasmids encoding HA-tagged human hnRNP H2 wild-type (WT) or the disease-associated variants p.R114W, p.R206Q, p.R206W or p.R206G, together with two knockdown sequences targeting endogenous rodent hnRNP H2. Data are expressed relative to control and presented as mean ± SEM from 3 independent neuronal preparations. At least 12–15 fields of view containing multiple dendrites analysed per condition in each experiment. Statistical significance was determined using two-tailed Mann–Whitney tests. p < 0.05 (*), p < 0.01 (**), p < 0.001 (***).

**Supplementary movie 1.**
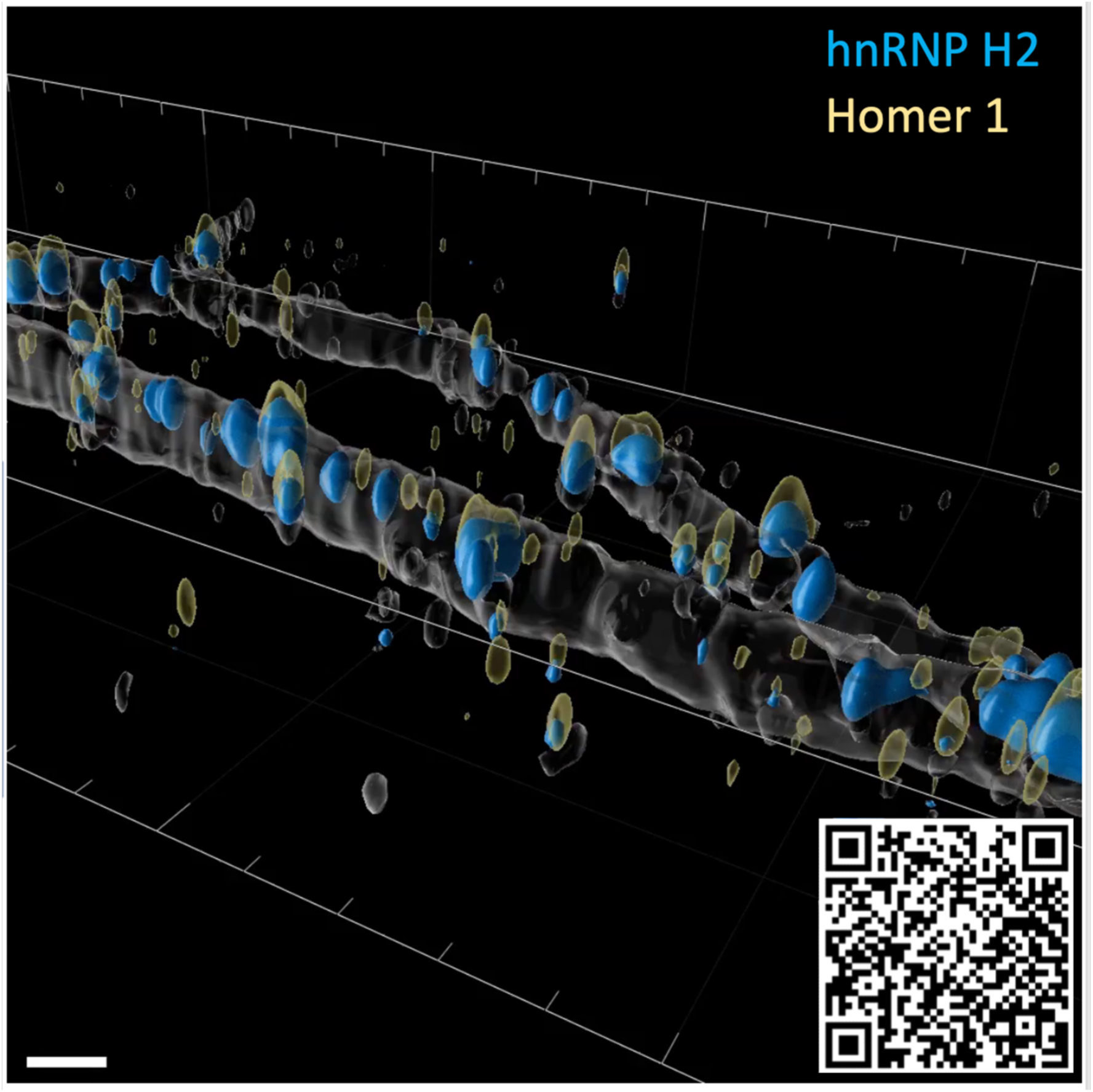
Three-dimensional visualization of hnRNP H2 localisation at excitatory synapses. Three-dimensional reconstruction of dendritic segments generated from Airyscan images and rendered using Imaris software. The movie shows hnRNP H2 puncta localised at the base of Homer1-positive dendritic spines, immediately below the postsynaptic marker Homer1. The corresponding still image is shown in Fig. 1c. Scale bar, 2 μm.

## Notes

### Competing Interest Statement

The authors have declared no competing interest.

